# SUMOylation is a Therapeutic Vulnerability in High-risk Neuroblastoma

**DOI:** 10.64898/2026.09.05.749647

**Authors:** Antti Kukkula, Saiganesh Sriraman, Verneri Virtanen, Sara L. M. Kuusela, Lara Kozina Bubnič, Ilkka Paatero, Vilja Pietiäinen, Daniel Bexell, Maria Sundvall

## Abstract

Neuroblastoma (NB) is one of the most common solid malignancies in children, and high-risk patients have poor prognosis, underscoring the need for novel therapeutic agents. The small ubiquitin-like modifier (SUMO) is a reversible post-translational modification that regulates various protein functions, and the first-in-class SUMOylation inhibitor TAK-981 (subasumstat) is currently in clinical trials in adult cancer patients. However, the role of SUMOylation in NB pathogenesis remains poorly characterized. We show that SUMOylation- catalyzing enzymes, *SAE1*, *SAE2*, and *UBE2I*, are highly expressed in malignant neuroendocrine cells, and their expression correlate with poor survival in NB patients and associate with advanced stage, *MYCN* amplifications, and downregulation of late differentiation markers. The peripheral nervous system lineage and NB cell lines are highly sensitive to the CRISPR knockout of *SAE1* and *SAE2*, suggesting SUMOylation as promising therapeutic target in NB. Accordingly, TAK-981-mediated inhibition of SUMOylation reduces growth of NB cells in both *in vitro* and *in vivo* models through induction of apoptosis and perturbation of differentiation-associated pathways. Sensitivity to SUMOylation inhibition is greatest in NB cells with low expression of favorable late neuroblast markers and is independent of *MYCN* amplification status. Moreover, TAK-981 is effective in combination with differentiation-inducing all-trans retinoic acid and the DNA methyltransferase inhibitor decitabine. Mechanistically, combination of TAK-981 and retinoic acid potently downregulates retinoic acid receptor alpha (RARα) expression, whereas inhibition of SUMOylation combined with decitabine induces a strong accumulation of DNA damage. Taken together, these findings establish SUMO pathway components as potential prognostic markers and SUMOylation as a therapeutic vulnerability in high-risk NB.

## Introduction

Neuroblastoma (NB) is the most common extracranial solid tumor in children and the most frequent malignancy during the first year of life(*1*). The median age at diagnosis is approximately 19 months, with 90% of cases occurring before 5 years of age(*1*). NB originates from developing neural crest cells, typically forming tumors in the adrenal glands or sympathetic ganglia. Its biological and clinical presentation is highly heterogeneous, ranging from spontaneously regressing tumors to widely disseminated aggressive metastatic disease. Patients are stratified into low-, intermediate-, and high-risk groups based on disease stage, histopathology, age at diagnosis, *MYCN* amplification status, and tumor cell ploidy or DNA index(*1–3*). Treatment strategies vary according to risk: low-risk patients may be managed with observation alone or surgical resection with or without low-intensity chemotherapy, whereas intermediate-risk patients receive moderate-intensity chemotherapy in combination with surgery(*1*, *4*). High-risk patients require intensive multimodal therapies, often including high-dose chemotherapy, surgery, radiotherapy, autologous stem cell transplantation, differentiation therapy with 13-cis-retinoic acid, and immunotherapy with anti-disialoganglioside (GD2) antibodies(*1*, *4*). While 5-year survival rates are excellent for low- and intermediate-risk NB patients (>95% and 90–95%, respectively), outcomes for high-risk patients remain poor (50– 60%)(*4*), highlighting the need for better prognostic markers and treatments for high-risk NB.

Germline mutations of paired-like homeobox 2B (*PHOX2B*) and anaplastic lymphoma kinase (*ALK*) have been linked to familial cases of NB(*5–8*), suggesting the involvement of sympathetic neuronal differentiation-associated and classical oncogenic receptor tyrosine kinase pathways in NB pathogenesis. In sporadic NB, the most common genetic alterations include somatic amplification of *MYCN*, found in approximately 20% of tumors, as well as somatic activating mutations and amplifications of *ALK*, which occur in 6–17% and 2–10% of cases, respectively (*7–12*). Other less frequent somatic alterations include inactivating mutations of *ATRX* and activating mutations of *PTPN11* and *NRAS*(*11*). While precision oncology in NB is still evolving, preclinical studies have identified potential strategies to target MYCN, and ALK inhibitors have shown promise in clinical trials of high-risk patients with genomic *ALK* alterations (*4*, *13–16*).

Dysregulation of post-translational modifications (PTMs) mediated by small proteins contributes to the pathogenesis of various cancers, and therefore the components regulating the PTMs are emerging as potential therapeutic targets and prognostic markers(*17*, *18*). SUMOylation is a reversible PTM in which small ubiquitin-like modifiers (SUMOs) are conjugated to substrate proteins, affecting their protein-protein interactions, stability, localization, and enzymatic activity(*19–21*). The SUMO pathway is regulated by SUMO E1- activating enzymes (SAE1/SAE2), the sole E2-conjugating enzyme UBC9 (encoded by *UBE2I*), and E3 ligases such as PIAS family proteins. DeSUMOylation and maturation of SUMO are mediated by sentrin-specific proteases (SENPs). Notably, several tumor suppressors and oncoproteins, including c-MYC and MYCN, are SUMO substrates(*17*, *18*, *22*, *23*). While previous data suggests that upregulation of SUMOylation is essential for the pathogenesis of certain c-MYC-dependent cancers, its association with *MYCN*-amplified NB remains unknown(*24–27*). The recent emergence of TAK-981/subasumstat (SAE inhibitor), the first SUMOylation-targeting inhibitor to enter clinical trials in cancer patients, has opened new opportunities for SUMOylation research due to its high specificity and potency(*17*, *18*, *28*).

Here, we show that the key SUMOylation-promoting factors, *SAE1*, *SAE2*, and *UBE2I*, are elevated in high-risk NB, particularly in *MYCN*-amplified or metastatic tumors, and correlate with poor patient survival. Experimentally, TAK-981 reduces the global conjugation of SUMOs to substrates and efficiently reduces the growth of multiple NB cell lines and 3D tumoroids irrespective of *MYCN*-amplification status, shows *in vivo* efficacy in a zebrafish NB xenograft model, and perturbs 3D growth in a NB patient-derived xenograft (PDX) model. We also show that NB cells expressing low levels of late neuroblast markers are more sensitive to SUMOylation inhibition. Mechanistically, inhibition of SUMOylation perturbs differentiation-associated pathways in NB and induces apoptosis. In addition, TAK-981 potentiates the effect of the differentiation-inducing drug all-trans retinoic acid (ATRA) in NB cells and is synergistic with DNA methyltransferase (DNMT) inhibitor decitabine. Collectively, our findings identify SUMOylation as a novel therapeutically targetable vulnerability in aggressive NB.

## Materials and methods

### Analysis of mRNA expression levels in cohorts and datasets

Survival analysis was performed using the Kocak (GSE45547)(*29*), SEQC (GSE62564)(*30*), Versteeg (GSE16476)(*31*), and Cangelosi(*32*) cohorts. Data overlapping with the SEQC cohort in the integrated Cangelosi cohort was removed from the analysis. Patients with available overall survival (OS) data (Kocak, n=476/649; SEQC, n=498/498; Cangelosi, n=417/417; Versteeg n=88/88) were stratified into high and low expression groups using median mRNA expression as the cutoff value for the respective Kaplan-Meier analyses.

Expression of SUMOylation regulators in different stage, *MYCN* amplification, and diagnosis age groups were analyzed from the Kocak cohort (n=649) and association with histopathology from the Therapeutically Applicable Research to Generate Effective Treatments (TARGET) cohort (n=220). The integrated dataset of NBAtlas(*33*) was utilized in the analyses of single-cell transcriptomic data. Expression data of NB murine models was analyzed from the GSE32386(*34*) dataset. Differences in expression levels of SUMOylation- promoting factors between ganglioneuroma (n=3), ganglioneuroblastoma (n=8 samples from 6 patients), and NB (53 samples from 52 patients) tumors was analyzed from GSE12460(*7*). Ganglioneuroma and ganglioneuroblastoma samples were pooled due to low number of samples. The effect of ATRA on expression of SUMOylation-promoting factors in NB cell lines was analyzed from GSE155000(*35*) and GSE9169(*36*). Survival and mRNA expression data (log2) were downloaded from R2: Genomics Analysis and Visualization Platform at http://r2.amc.nl, except for the histopathology-related mRNA expression data, which were acquired from cBioPortal at https://www.cbioportal.org/.

Gene expression data of ATRA- and DMSO-treated CHP-134 cells(*37*) of GSE253785 dataset were downloaded from GEO2R at https://www.ncbi.nlm.nih.gov/geo/geo2r/. Volcano plot of differentially expressed genes was generated with VolcaNoseR at https://huygens.science.uva.nl/VolcaNoseR/ using an FDR/adjusted p-value <0.001 (Benjamini–Hochberg) and log2 fold change >0.5 as the threshold for statistical significance.

### Analysis of genetic alterations

Copy number alterations of SUMO machinery components in NB patients and cell lines was analyzed from the TARGET cohort (n=59) and the Cancer Cell Line Encyclopedia (CCLE), respectively. TARGET data was downloaded from cBioPortal and CCLE data was downloaded from cBioPortal and Dependency Map (DepMap) portal at https://depmap.org/portal/.

### Putative MYCN transcriptional targets

UCSC Genome Browser was used to examine the ENCODE candidate promoter-like signatures and enhancer-like signatures of ReMap ChIP-seq tracks for MYCN in *SAE1*, *SAE2*, and *UBE2I*. The graphs visualizing observed peaks in the putative promoter and enhancer sites were downloaded from http://genome.ucsc.edu.

### Gene set overlap analysis

Data of genes co-expressing with *SAE1*, *SAE2*, *UBE2I*, and *SUMO1*–*3* were downloaded from https://r2.amc.nl/ for the Kocak, SEQC, Versteeg, and Cangelosi cohorts. Data overlapping with the SEQC cohort in the Cangelosi cohort was excluded from the analysis. Lists of commonly correlating genes were generated by identifying the genes featured among the top 1000 positively (FDR<0.05) correlated genes in all cohorts. The common genes were compared to Molecular Signatures Database (MSigDB) (version 7.5) Gene Ontology: Biological Processes (GO:BP) and Hallmark gene sets and significant overlaps were identified.

### GSEA

For GSEA (version 4.3.2) of the Kocak cohort, genes were pre-ranked based on Pearson correlation of *SAE1* co-expressed genes. Genes for GSEA of mouse tissues of NB models (GSE32386) and ATRA-treated CHP-134 cells (GSE253785) were pre-ranked by log2 fold change x –log10 adjusted p value. Volcano plot of Reactome gene sets that were upregulated and downregulated in NB mouse tissues compared to normal adrenal medulla tissues was generated in VolcaNoseR. Gene sets with FDR<0.05 and normalized enrichment score (NES) >1 or <-1 were considered statistically significant.

### Analysis of CRISPR gene effects in Oncotree lineages and NB cell lines

Gene effects of *SAE1*, *SAE2*, and *UBE2I* were analyzed from the CRISPR (DepMap Public 26Q1+Score, Chronos) dataset which contains CRISPR sensitivity data of cell lines (n=1208) derived from various different cancer types and lineages. The data was downloaded from https://depmap.org/portal/.

### Cell culture

*MYCN*-non-amplified SH-SY5Y and GI-ME-N as well as, *MYCN*-amplified CHP-134, KELLY, IMR-32, LAN-1, and SK-N-BE(2) NB cell lines were purchased from Deutsche Sammlung von Mikroorganismen und Zellkulturen (DSMZ). We kindly thank Dr. Ruth Palmer for providing SK-N-AS (*MYCN*-non-amplified) cells. LU-NB-3 PDX-derived NB cells were obtained from Biobank Sweden with permission (2016/1055, SC2595). SH-SY5Y were cultured in DMEM (Gibco) supplemented with 20% heat-inactivated (h.i.) FBS (Gibco). GI-ME-N, CHP-134, and KELLY cells were cultured in RPMI (Gibco) supplemented with 10% h.i. FBS and 2 mM glutamine (Lonza). IMR-32 and LAN-1 were cultured in RPMI supplemented with 20% h.i. FBS, 2 mM glutamine and 1% non-essential amino acids (Gibco). SK-N-BE(2) were cultured in a 1:1 mixture of Ham’s F12 (Gibco) and EMEM (ATCC) supplemented with 10% h.i. FBS. SK-N-AS were cultured in DMEM supplemented with 10% h.i. FBS, 1% non-essential amino acids, and 1% sodium pyruvate (Gibco). LU-NB-3 PDX cells were grown in stem cell medium consisting of 1:4 mixture of Ham’s F-12 and low glucose DMEM (#21885025, Gibco) supplemented with 40 ng/ml FGF2 (#100-18B, PeproTech/Gibco), 20 ng/ml EGF (#AF-100-15, PeproTech/Gibco), B-27 supplement minus vitamin A (#12587010, Gibco), and 1% pen-strep. Cell lines were cultured at 37 °C (5% CO2) and regularly tested to be mycoplasma-negative.

### MTS and WST-8 cell viability assays

Cells were cultured as triplicates in 96-well plates in monolayer drug experiments, and cell viability was determined with CellTiter 96® AQueous Non-Radioactive Cell Proliferation MTS Assay (#G5430; Promega) by measuring the absorbance at 490 nm. LU-NB-3 3D spheroids were cultured as duplicates in 96-well plates and cell viability was determined with Cell Counting Kit 8 (WST-8) (ab228554, Abcam) at 450 nm. Absorbance was measured with Wallac Victor2 1420 Multilabel Counter (PerkinElmer).

### 3D growth assays

Cells were reconstituted in 25% basement membrane matrix (Corning; Matrigel® Growth Factor Reduced or R&D Systems; Cultrex Reduced Growth Factor Basement Membrane Extract) and seeded in 96-well plates precoated with 50% basement membrane matrix (Matrigel or Cultrex). DMSO or TAK-981 (0.1 and 0.5 µM) diluted in 150 µl of medium was added on top of 40 µl cell layer and 30 µl precoat after 1-hour incubation at 37 °C. Medium was replaced every 2–3 days with fresh medium containing treatments. Duplicate wells were imaged on day 10 for a minimum of three 4x brightfield images/well using EVOS M5000 (Thermo Fisher Scientific). Spheroid size was measured in ImageJ with segmenting by manual thresholding and manual exclusion of spheroids merged due to high initial proximity.

### Drug sensitivity and resistance testing for synergistic interactions

Drug sensitivity and resistance testing (DSRT) of IMR-32 cells was performed using barcoded 384-well u-bottom drug plates (Corning) prepared by the FIMM High Throughput Biomedicine unit (FIMM, HiLIFE, University of Helsinki, Finland). Each plate contained a matrix of 7- point concentration series of single agents and drug combinations. Drugs were pre-dispensed using acoustic liquid handling (Echo 550, Labcyte), and benzoyl chloride (BzCl) and DMSO were used as internal positive and negative controls, respectively. The cell viability was measured using the CellTiter-Glo® Luminescent Cell Viability Assay (Promega), measuring the intracellular ATP, and luminescence was read on a PHERAstar FS plate reader (BMG Labtech). Viability data were normalized to DMSO controls, and synergy analysis was performed using SynergyFinder 2.0(*38*) at https://synergyfinder.fimm.fi.

### Cell lysis and western blot analysis

Cells lysis and western blot were performed as previously described(*39*), and used antibodies are listed in Supplementary table S1. Densitometry was done with Image Lab software (version 6.0.1, Bio-Rad).

### Immunofluorescence staining

Cells were cultured on coverslips in 24-well plates and subjected to DMSO or TAK-981 (0.5 µM) for 48 hours. Immunofluorescence staining of SUMO1 (ab133352, 1:250; Abcam) and SUMO2/3 (ab81371, 1:2000; Abcam) was performed using anti-rabbit AlexaFluor 488 (#A11034) and anti-mouse AlexaFluor 555 (#A21422) secondary antibodies as previously described(*39*). The samples were imaged with the Eclipse Ni microscope (Nikon) equipped with the DS-Qi2 camera (Nikon). Images were processed in ImageJ.

### Zebrafish embryo NB xenografts

Zebrafish experiments were carried out under the license ESAVI/38339/2024, granted by the Project Authorization Board of the Regional State Administrative Agency for Southern Finland in accordance with the regulations of the Finnish Act on Animal Experimentation (62/2006) in the Zebrafish Core Facility of the Turku Bioscience Centre. The study was carried out in compliance with the ARRIVE guidelines.

Wild type AB zebrafish embryos were obtained using natural spawning in breeding tanks. The embryos were cultured in E3 + PTU medium at 28.5 °C until subjected to microinjection of CellTracker Green-labeled (#C7025, Invitrogen) IMR-32 cells using Nanoject II microinjector (Drummond Scientific) under a Zeiss AxioZoom.V16 fluorescence stereomicroscope (Zeiss). At 1-day post-injection (dpi), embryos were transferred into a 96-well glass-bottom imaging plate (1 embryo/well) in E3 + PTU + pen-strep (1:200) medium, and into 34 °C incubator. Formal randomization or blinding was not used. Embryos were treated with DMSO (n=24) or TAK-981 (1 μM; n=24) at 1 dpi. The embryos were anesthetized with 200 mg/l tricaine and imaged at 1 dpi to establish the baseline xenograft measurements, and at 4 dpi after the drug treatments, using a Nikon Ti2 widefield fluorescence microscope (Nikon). Primary tumor growth was analyzed from fluorescence images in ImageJ.

### Statistical analysis

Statistical analyses were performed using GraphPad Prism 10.1.2 software and R Studio (R Project for Statistical Computing). Welch’s or Student’s t-test were used for testing statistical significance between two groups. Log-rank test was used for testing statistical significance in survival analyses. Pearson correlation was used for determining statistically significant co- expression between genes or proteins. Kruskal-Wallis test was used to compare mRNA expression levels across multiple groups in the scRNA-seq data.

## Results

### High expression of SUMOylation-promoting factors associates with poor prognosis and high-risk features in NB patients

First, we analyzed the prognostic impact of SUMO machinery components in the Kocak(*29*), SEQC(*30*), Versteeg(*31*), and Cangelosi(*32*) NB patient cohorts (Supplementary Fig. S1A). Strikingly, high mRNA expression of the major SUMOylation-promoting factors *SAE1* (Kocak p=5.8e−8, SEQC p=4.8e−15, Cangelosi p=1.2e−8, Versteeg p=3.6e−5), *SAE2* (Kocak p=3.9e−7, SEQC p=2.4e−17, Cangelosi p=5.0e−3, Versteeg p=7.2e−6), and *UBE2I* (Kocak p=1.7e−7, SEQC p=7.2e−10, Cangelosi p=3.8e−9, Versteeg p=4.1e−4) consistently associated with poor overall survival (OS) in all of the cohorts (Fig. 1A and Supplementary Fig. S1A) indicating a strong link between increased SUMOylation activity and adverse clinical outcome. We next examined the prognostic relevance of SUMO-specific proteases. Expression of *SENP2*, *SENP6*, and *SENP7* correlated with OS in all cohorts. Notably, high *SENP2* expression was associated with poor OS (Kocak p=7.0e−10, SEQC p=4.3e−5, Cangelosi p=1.6e−7, Versteeg p=2.7e−3), whereas high expression levels of *SENP6* (Kocak p=7.2e−03, SEQC p=1.3e−6, Cangelosi p=3.5e−6, Versteeg p=2.1e−6) and *SENP7* (Kocak p=4.8e−11, SEQC 1.7e−10, Cangelosi p=1.8e−14, Versteeg p=9.8e−7) correlated with good prognosis. Overall, these findings suggest that SUMOylation-promoting factors correlate more uniformly with poor survival, whereas individual deSUMOylases may exert distinct, context-dependent roles in NB.

**Figure 1.**
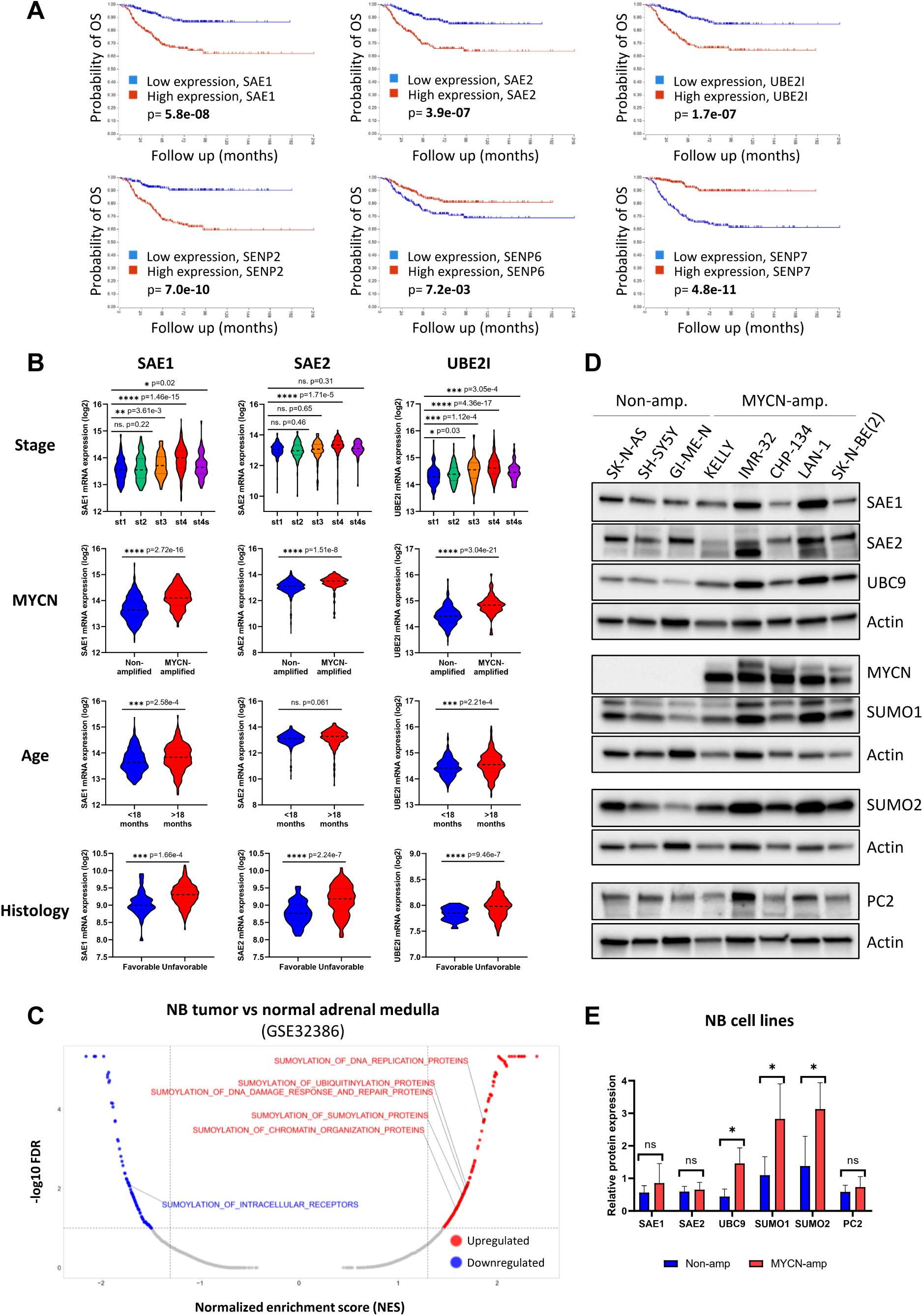
SUMOylation-regulating factors have prognostic significance and associate with high-risk clinical features in NB patients. **(A)** Representative Kaplan-Meier plots of *SAE1*, *SAE2*, *UBE2I*, *SENP2*, *SENP6*, and *SENP7* impact on OS of NB patients in the Kocak cohort (n=476). Median mRNA expression value was used as a cutoff for group stratification. Statistical significance was determined with the log-rank test. **(B)** Violin plots depicting the association of *SAE1*, *SAE2*, and *UBE2I* expression with low- and high-risk clinical features in NB patients. Correlation with disease stage (st) was analyzed from patients with st1 (n=153), st2 (n=113), st3 (n=91), st4 (n=214), and st4s (n=78) tumors. Association with *MYCN* amplification status was analyzed between *MYCN*-non-amplified (n=550) and *MYCN*- amplified (n=93) tumors. Correlation with diagnosis age was analyzed from patients who were <18 months (n=414) and >18 months (n=235) of age. Correlation with histopathological prognosis was evaluated from tumors with favorable (n=37) and unfavorable (n=183) histology. Association with stage, *MYCN* status, and age were analyzed from the Kocak cohort and correlation with histology was analyzed from the TARGET cohort. Welch’s t-test was used for the statistical testing (* p < 0.05, ** p < 0.01, *** p < 0.001, **** p < 0.0001). **(C)** Volcano plot depicting upregulated and downregulated Reactome gene sets in mouse NB tumor (n=10) tissues compared to normal adrenal medulla (n=3) tissues, based on GSEA analysis. Positively and negatively enriched SUMOylation-related gene sets are highlighted in red and blue, respectively (FDR <0.05, normalized enrichment score, NES >1 or <-1). Tumor samples include *ALK*- (n=3), *MYCN*- (n=4), and *ALK*+*MYCN*-driven (n=3) tumors. Analysis was done from the GSE32386 dataset. **(D)** Western blot analysis of SAE1, SAE2, UBC9, MYCN, SUMO1, SUMO2, and PC2 protein levels in human NB cell lines and **(E)** a bar graph comparing the protein levels between *MYCN*-non-amplified and *MYCN*-amplified cell lines using actin as the loading control. Student’s t-test was used for the statistical testing (* p < 0.05).

Next, we analyzed genetic alterations affecting SUMO pathway components from the TARGET(*11*) cohort together with NB cell line data from the CCLE (Supplementary Fig. S1B and C). Amplification of the SUMO E3 ligase *PC2* (10.2%, n=6/59) and *SUMO2* (8.5%, n=5/59) were the most common alterations in patients, whereas amplifications of *UBE2I* (1.7%, n=1/59) and *PIAS2* (1.7%, n=1/59) occurred only in single cases. Deletions were rare and limited to isolate events in *SUMO1* (1.7%, n=1/59) and *RSUME* (1.7%, n=1/59). In line with patient data, amplifications of *PC2* (35.3%, n=6/17) and *SUMO2* (29.4%, n=5/17) were also the most frequent alterations in NB cell lines. Other alterations in CCLE data included amplifications of *PIAS3* (11.8%) and deletions of *PIAS2* (23.5%), *SENP3* (23.5%), *RSUME* (17.3%), *RANBP2* (11.8%), and *SAE1* (11.8%). Interestingly, we identified a missense mutation of *SAE1* (K45E) in SH-SY5Y cells and its parental cell line SK-N-SH, while no other mutations of putative functional impact were observed. Overall, genetic alterations of SUMOylation-regulating factors were relatively common in NB and, together with the survival associations, support a functional role for SUMO pathway activity in NB pathogenesis.

To improve our understanding of the role of SUMO pathway in NB, we analyzed the mRNA expression of SUMO machinery components across key clinical variables. First, we analyzed SUMOylation-promoting factors with the strongest significance in the survival analyses (Fig. 1B). High expression of *SAE1* correlated with advanced disease stage (stage 1 vs. 4, p=1.46e−15), higher number of *MYCN* amplifications (p=2.72e−16), and older age at diagnosis (>18 months, p=2.58e−4). Correspondingly, *SAE2* expression associated with advanced stage (p=1.71e−5) and *MYCN* amplifications (p=1.51e−8) but did not significantly correlate with patient age (p=0.061). *UBE2I* expression showed a strong positive correlation with advanced stage (p=4.36e−17), *MYCN* amplifications (p=3.04e−21), and older age (p=2.21e−4) in patients. Moreover, high expression of *SAE1* (p=1.656e−4), *SAE2* (p=2.24e−7), and *UBE2I* (p=9.46e−7) was associated with unfavorable histology.

Similarly, *SENP2*, *SENP6*, and *SENP7* were associated with clinical features that align with their prognostic directions (Supplementary Fig. S2A–C). High expression of *SENP2* correlated with advanced stage (p=3.62e−20), *MYCN* amplifications (p=8.95e−12), and older age (p=2.38e−11) but not with unfavorable histology (p=0.830). In contrast, high expression of *SENP6* and *SENP7* was associated with lower disease stage (p=7.25e−10 and p=5.77e−15, respectively), lower frequency of *MYCN* amplifications (p= 2.64e−10 and p= 5.92e−11), and younger age at diagnosis (p=9.15e−8 and p=5.01e−25). High expression of *SENP6* also associated with favorable histology (p=1.442e−3), and *SENP7* displayed a similar trend (p=0.083). Several additional SUMO regulators were likewise associated with favorable or aggressive clinical features, suggesting a more extensive role for the SUMO pathway in driving NB aggressiveness. (Supplementary Fig. S2D).

To determine whether SUMO pathway activation occurs during tumor development, we analyzed RNA-seq data from previously published transgenic NB mouse models(*34*). Mouse NB tumor tissues revealed upregulation of several SUMOylation-related gene sets (Fig. 1C, Supplementary table S2) as well as higher expression of *Sae1*, *Sae2*, *Ube2i*, and *Sumo1*–*3* (Supplementary Fig. S2E) compared with normal adrenal medulla samples, proposing that SUMOylation is elevated during NB tumorigenesis. *MYCN*-driven transgenic NB mice tumors showed higher expression of SUMOylation-promoting factors than *ALK*-driven tumors, in line with our findings in patient samples.

We evaluated the protein levels of SAE1, SAE2, UBC9, SUMO1, SUMO2, and PC2 in a panel of *MYCN*-non-amplified and *MYCN*-amplified human NB cell lines by western blot (Fig. 1D). While SAE1, SAE2, and PC2 expression varied across cell lines, the highest levels were detected in the *MYCN*-amplified LAN-1 and IMR-32 cell lines. Interestingly, the protein levels of UBC9, SUMO1, and SUMO2 were higher in *MYCN*-amplified cell lines compared with non-amplified cell lines (Fig. 1E). Given the high expression of these factors in *MYCN*- amplified NBs, we investigated whether *MYCN* could contribute to their upregulation. By utilizing the UCSC Genome Browser to examine the ReMap ChIP-seq tracks for MYCN, we observed peaks in the putative promoter and enhancer sites of *SAE1*, *SAE2*, and *UBE2I*, suggesting potential transcriptional regulation by MYCN (Supplementary Fig. S3A–C). As NB patients harboring *MYCN* amplifications generally have poorer prognosis than patients without *MYCN* amplification, we also evaluated the survival impact of *SAE1*, *SAE2*, and *UBE2I* separately in non-amplified patients, and observed that they retain their prognostic value in the non-amplified subgroup, suggesting that their prognostic effect is not secondary to putative upregulation by amplified *MYCN* (Supplementary Fig. S3D).

### Expression of *SAE1*, *SAE2*, and *UBE2I* associate with poor differentiation status of neuroblasts

As SUMOylation-promoting factors were significantly correlating with poor prognosis and aggressive clinical features, they were further analysed by examination of single-cell transcriptomic data of the NBAtlas. Our analysis highlighted that expression levels of *SAE1*, *SAE2*, and *UBE2I* are enriched in neuroendocrine/cancer cells compared to other cell types of the tumor microenvironment (Fig. 2A).

**Figure 2.**
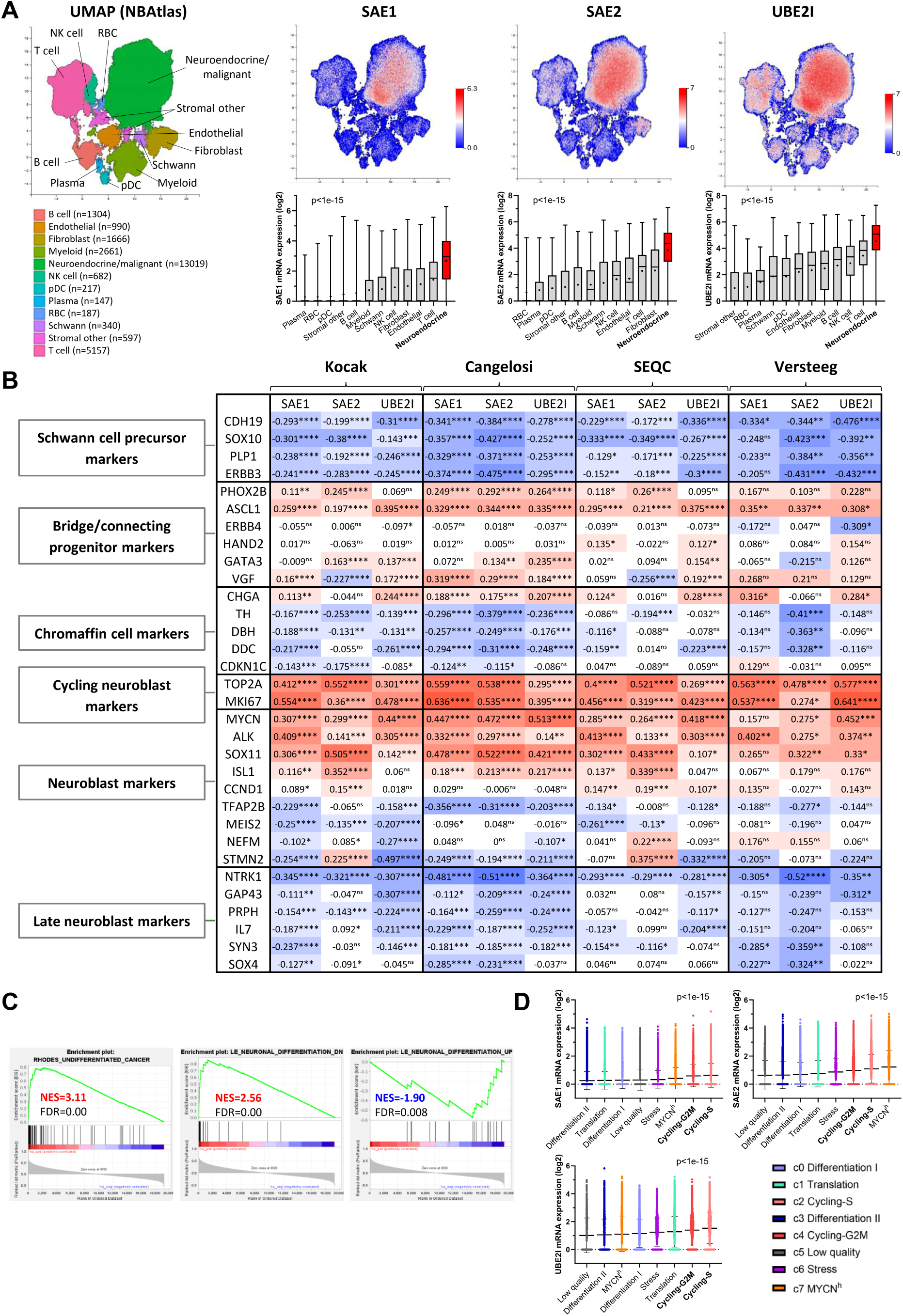
*SAE1*, *SAE2*, and *UBE2I* are enriched in NB cancer cells and associate with poor differentiation status of neuroblasts. **(A**) UMAP of NBAtlas (n=126872) colored according to cell type and UMAPs of *SAE1*, *SAE2*, and *UBE2I* expression in NBAtlas and box plots of expression levels in different cell types of the tumor microenvironment. Statistical significance was determined with Kruskal-Wallis test. **(B)** Heatmap representing co-expression of *SAE1*, *SAE2*, and *UBE2I* with markers of neural crest-derived cell differentiation trajectories in the Kocak, Cangelosi, SEQC, and Versteeg cohorts. Red and blue indicate positive and negative co-expression, respectively. Statistical significance was determined using Pearson correlation analysis (* p < 0.05, ** p < 0.01, *** p < 0.001, **** p < 0.0001). **(C)** GSEA analysis of *SAE1* in Kocak cohort for RHODES_UNDIFFERENTIATED_CANCER (metasignature commonly upregulated in undifferentiated cancers) and LE_NEURONAL_DIFFERENTIATION_UP/DN gene sets (genes upregulated or downregulated during neuroblastoma cell differentiation). **(D)** Single-cell expression of *SAE1*, *SAE2*, and *UBE2I* in the neuroendocrine/cancer cell clusters of the NBAtlas dataset ranked from lowest to highest mean expression. Statistical significance was determined with Kruskal- Wallis test.

To examine which pathways and biological processes are associated with the SUMO pathway in NB, we analyzed patient tumor bulk mRNA expression datasets to determine the common transcripts co-expressed with SUMOylation-promoting factors between the cohorts (Supplementary table S3). These transcripts were characterized for overlaps with the GO:BP and Hallmark gene sets using MSigDB (Supplementary Fig. S4). Co-expressed transcripts of *SAE1*, *SAE2*, *UBE2I*, and *SUMO2* were strongly associated with various cell cycle-related gene sets, including E2F targets, G2/M checkpoint, and MYC targets, indicating a robust link to proliferative signaling. Beyond cell cycle-related processes, *SAE2* correlated with the biogenesis of ribonucleoproteins and ribosomes. *SUMO1* and *SUMO3* were associated with MYC targets, mTORC1 signaling, oxidative phosphorylation, and several biosynthetic processes, suggesting broader roles in growth and metabolic regulation.

Higher levels of SUMOylation-promoting factors were observed in NB tumors with unfavorable histopathology and other high-risk features, suggesting an association between increased SUMOylation and poor differentiation status. To further investigate this relationship, we analyzed *SAE1*, *SAE2*, and *UBE2I* expression in other neuroblastic tumors using a previously published dataset(*7*) and found higher expression in NB tumors compared with intermediately-matured ganglioneuroblastoma and well-matured ganglioneuroma tumors (Supplementary Fig. S5). To identify which cell populations are linked to high SUMOylation activity in NB tissues, we examined the co-expression of *SAE1*, *SAE2*, and *UBE2I* with neural crest cell differentiation markers(*40*) across the four NB cohorts (Fig. 2B). Expression of *SAE1*, *SAE2*, and *UBE2I* correlated negatively with the expression of Schwann cell precursor (SCP) markers *CDH19*, *SOX10*, and *ERBB3*, and chromaffin cell markers *TH*, *DBH*, and *DDC*, with the exception of positively co-expressed *CHGA*. In contrast, SUMOylation-promoting factors were positively associated with the bridge/connecting progenitor markers *PHOX2B* and *ASCL1*. Co-expression with additional bridge/connecting progenitor markers *GATA3* and *VGF* was heterogeneous but predominantly positive, whereas no clear correlation was observed with *ErbB4* or *HAND2*.

Notably, SUMOylation-promoting factors were positively associated with different neuroblast differentiation trajectories and risk markers. *SAE1*, *SAE2*, and *UBE2I* correlated positively with *MYCN*, *SOX11*, and *ISL1,* as well as the early neuroblast marker *ALK,* whereas correlation with *CCND1*, *TFAP2B*, *MEIS2*, *NEFM*, and *STMN2* levels was relatively heterogeneous across cohorts. Strong positive correlation was observed with cycling neuroblast markers *MKI67* and *TOP2A,* which characterize the actively proliferating subpopulation of NB cells associated with aggressive disease and poor prognosis(*40*). In contrast, SUMOylation- promoting factors had predominantly negative correlations with the expression of favorable late neuroblast markers *NTRK1*, *GAP43*, *PRPH*, *IL7*, *SYN3*, and *SOX4*. Overall, these findings indicate that SUMOylation-promoting factors mainly associate with elevated expression of markers associated with high-risk disease and poor differentiation state, while correlating negatively with favorable and late neuroblast state markers.

In accordance, GSEA analysis of *SAE1* showed strong enrichment for a transcriptional metasignature(*41*) commonly upregulated in various types of undifferentiated cancer (Fig. 2C). Moreover, *SAE1* had positive enrichment for a set of genes downregulated upon NB cell differentiation(*42*), while having negative enrichment for a gene set upregulated during differentiation. We also examined the single-cell expression of *SAE1*, *SAE2*, and *UBE2I* in the neuroendocrine/cancer cell subsets of the NBAtlas (Fig. 2D). Similarly to the bulk mRNA analyses, we observed higher expression of *SAE1*, *SAE2*, and *UBE2I* in the high-risk/aggressive cycling-S and cycling-G2M cell clusters when compared to the low-risk/favorable differentiation clusters I and II.

### NB cells with low expression of favorable late neuroblast markers are the most sensitive to SUMOylation inhibition

We analyzed CRISPR knockout data of various cell lines using the DepMap database and ranked the Oncotree tissue lineages based on the mean CRISPR gene effects of *SAE1*, *SAE2*, and *UBE2I* to determine which cell lineages are the most dependent on expression of SUMOylation-promoting factors. Peripheral nervous system (NB n=41; nerve sheath tumor n=6) was identified as the most sensitive lineage for *SAE1* knockout and ranked as the second most sensitive lineage for *SAE2* knockout (Fig. 3A), while *UBE2I* knockout ranked near the mean gene effect of all cell lines (Supplementary Fig. S6A). Analysis based on primary disease type identified NB and Ewing sarcoma as the only cancers that are sensitive for both *SAE1* (NB, p=3.28e-3) and *SAE2* (NB, p=9.22e-3) knockout when compared to all other cell lines (Supplementary Fig. S6B). These data suggest that expression of *SAE1* and *SAE2* contributes to the growth of peripheral nervous system-derived cancers, making them promising drug targets in NB.

**Figure 3.**
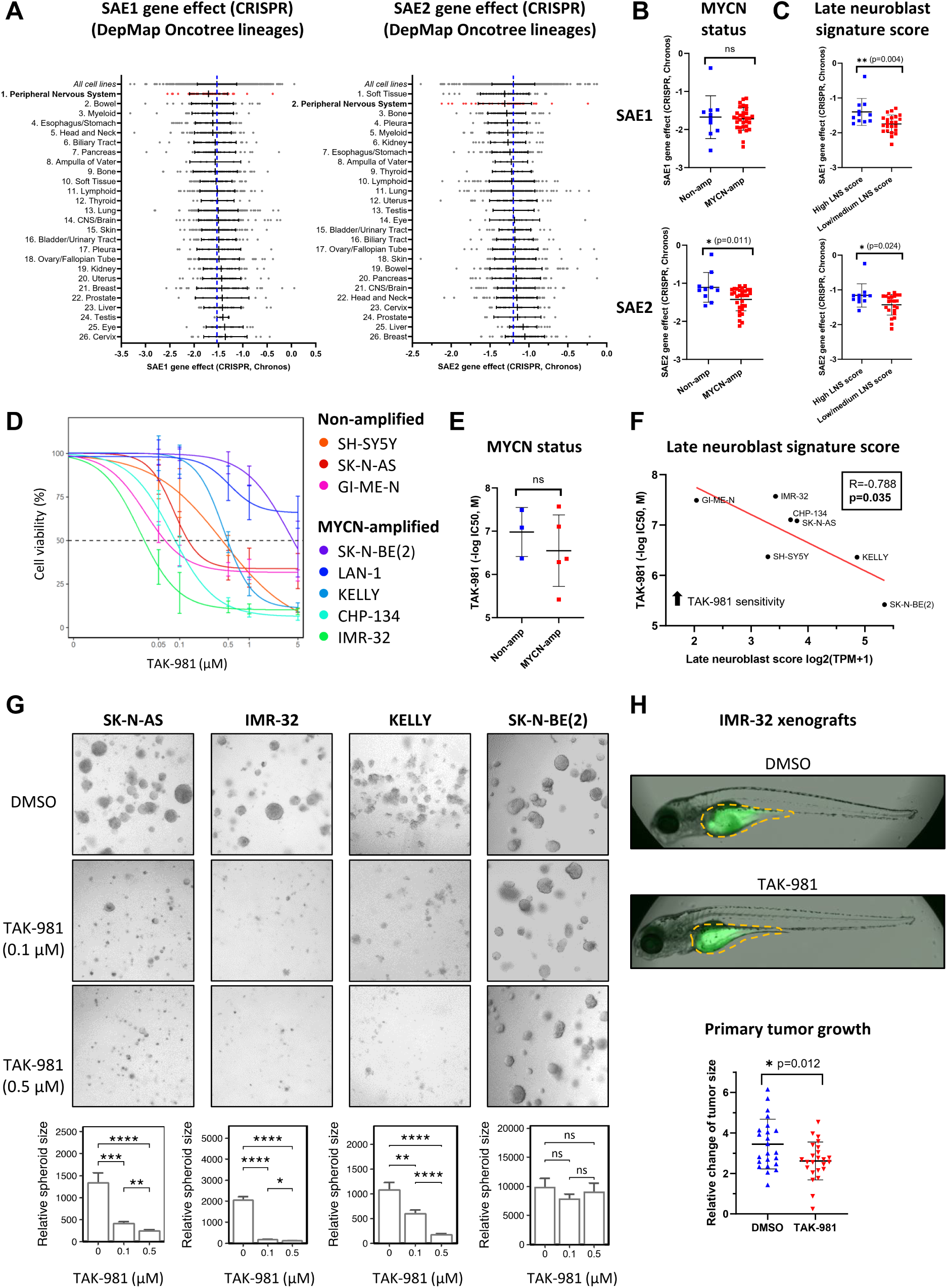
SAE inhibitor TAK-981 shows in vitro and in vivo efficacy in NB models. **(A)** DepMap Oncotree lineages ranked based on the mean gene effects of *SAE1* or *SAE2* analyzed from the CRISPR (DepMap Public 26Q1+Score, Chronos) dataset (n=1208). A lower gene effect score indicates a higher likelihood that the gene is important for cell growth/viability. The blue line indicates the mean gene effect across all cell lines. **(B)** *SAE1* gene effect in *MYCN*-non-amplified (n=10) and *MYCN*-amplified (n=31) NB cell lines tested with Student’s t-test. **(C)** *SAE1* gene effect in NB cell lines expressing high (top tertile) and low/medium (bottom tertiles) late neuroblast score tested with Student’s t-test. **(D)** Cell viability of *MYCN*- non-amplified (GI-ME-N, SK-N-AS, and SH-SY5Y) and *MYCN*-amplified (IMR-32, CHP- 134, KELLY, LAN-1, and SK-N-BE(2)) NB cell lines treated with TAK-981 for 96 hours (n=3). **(E)** Comparison of TAK-981 IC50 values (-log, M) in MYCN-non-amplified (n=3) and MYCN-amplified (n=5) NB cell lines tested with Student’s t-test. **(F)** Pearson correlation analysis of TAK-981 IC50 values (-log, M) and late neuroblast score in NB cell lines. **(G)** Representative brightfield images of non-amplified SK-N-AS cells, and *MYCN*-amplified IMR-32, KELLY, and SK-N-BE(2) cells treated with DMSO and TAK-981 (0.1 and 0.5 µM) grown in Matrigel (day 10). Bar graphs with SEM indicate the relative spheroid size (SK-N- AS, IMR-32, KELLY, and SK-N-BE(2)). Student’s t-test was used for statistical testing (* p < 0.05, ** p < 0.01, *** p < 0.001, **** p < 0.0001). Number of spheroids: SK-N-AS (DMSO n=65; TAK-981 0.1 µM n=103; TAK-981 0.5 µM n=80), IMR-32 (DMSO n=134; TAK-981 0.1 n=168; TAK-981 0.5 µM n=134), KELLY (DMSO n=40; TAK-981 0.1 µM n=63; TAK- 981 0.5 µM n=122), and SK-N-BE(2) (DMSO n=20; TAK-981 0.1 µM n=42; TAK-981 0.5 n=24). **(H)** Representative merged brightfield and fluorescent images of zebrafish embryos microinjected with CellTracker Green-labeled IMR-32 cells into the yolk sac 4 days after injection. The scatter plot displays the relative primary tumor growth (size 4 days after injection / size 1 day after injection) in DMSO (n=23) and TAK-981 (1 μM, n=23) groups after 72 hours of treatment. The tumor area was measured from the fluorescence images, and Student’s t-test was used for statistical testing (* p < 0.05).

Comparison of *SAE1* and *SAE2* CRISPR sensitivity between *MYCN*-non-amplified and *MYCN*-amplified NB cell lines showed no difference in *SAE1* knockout sensitivity, whereas *MYCN*-amplified cell lines were more sensitive for *SAE2* knockout (Fig. 3B). As the SUMO-activating enzyme requires the heterodimerization of SAE1 and SAE2 to be functional, we also analyzed the combined mean gene effect of *SAE1* and *SAE2* in *MYCN*-non-amplified and *MYCN*-amplified cell lines, which showed no significant difference in sensitivity based of *MYCN* amplification status (Supplementary Fig. S6C). Similarly, *ALK* mutation/amplification status had no impact on *SAE1* or *SAE2* CRISPR sensitivity (Supplementary Fig. S6D). Due to our observations linking NB cell differentiation and SUMOylation, we wanted to determine if the differentiation status of the NB cells affects their sensitivity to knockout of SUMOylation- promoting factors. We determined a late neuroblast signature score comprised of the combined mean expression of the favorable late neuroblast markers(*40*) *NTRK1*, *GAP43*, *PRPH*, *IL7*, *SYN3*, and *SOX4* for each NB cell line available in DepMap (Supplementary Fig. S6E). Interestingly, NB cell lines expressing low/medium late neuroblast score were more sensitive for *SAE1* and *SAE2* knockout (Fig. 3C), as well as the combined mean gene effect of *SAE1* and *SAE2* knockout (Supplementary Fig. S6F), which may indicate that less differentiated NB cells are more sensitive for SUMOylation inhibition.

Based on these findings, we evaluated the effect of pharmacological SUMOylation inhibition on the growth of NB cells using the first-in-class SAE inhibitor TAK-981 which forms an adduct with SUMO as the inhibitory species within the enzyme catalytic site of the SAE1/SAE2 heterodimer(*28*). TAK-981 induced a concentration-dependent reduction of cell viability in *MYCN*-non-amplified GI-ME-N, SK-N-AS, and SH-SY5Y cells, as well as in *MYCN*-amplified IMR-32, CHP-134, KELLY, LAN-1, and SK-N-BE(2) cells in 2D monolayer assay after 96 hours of treatment (Fig. 3D). Notable differences in sensitivity to TAK-981 were detected between the cell lines, ranging from highly sensitive IMR-32 and GI-ME-N to more resistant SK-N-BE(2) and LAN-1. No clear difference in sensitivity to TAK-981 was observed between *MYCN*-amplified and non-amplified cell lines (Fig. 3E) or *ALK* wild type and *ALK*- mutated/-amplified cell lines (Supplementary Fig. S6G). Consistent with DepMap CRISPR data, lower late neuroblast signature score was significantly correlated with higher sensitivity to TAK-981 (Fig. 3F). Interestingly, two of the TAK-981-sensitive cell lines (IMR-32 and SK-N-AS) also harbored *SUMO2* and *PC2* amplifications; however, copy number alteration status data was not available for all cell lines. Next, we evaluated the effect of TAK-981 on 3D- cultured NB models, which more closely resemble *in vivo* tissue conditions. TAK-981 reduced spheroid formation and growth in *MYCN*-non-amplified SK-N-AS, *MYCN*-amplified IMR-32, and KELLY cells, whereas it did not significantly affect the growth of SK-N-BE(2) cells (Fig. 3G). TAK-981 also reduced growth of non-amplified GI-ME-N cells which did not grow as classical spheroids but formed branched cell structures in 3D (Supplementary Fig. S6H). Finally, we confirmed the *in vivo* efficacy of TAK-981 using a zebrafish IMR-32 xenograft model, in which TAK-981 reduced the growth of primary tumors (p=0.012) in zebrafish embryos after 72 hours of treatment.

Taken together, TAK-981 is effective across multiple NB models. Less differentiated NB cells are more sensitive to inhibition of SUMOylation regardless of their *MYCN* or *ALK* alteration status. However, the impact of genetic alterations of SUMO pathway regulators on sensitivity of cancer cells to SUMOylation inhibition requires further validation.

### Inhibition of global SUMOylation induces apoptosis and partial differentiation in NB cells

To investigate the molecular effect of TAK-981, we analyzed the conjugation levels of SUMO1 and SUMO2 by western blot after 48-hour treatment. TAK-981 treatment resulted in a concentration-dependent reduction in global conjugation of SUMO1 and SUMO2, accompanied by increasing levels of free SUMO1 and SUMO2 in *MYCN*-non-amplified SK-N-AS and *MYCN*-amplified KELLY, IMR-32, and LAN-1 cells with varying sensitivity to TAK-981, suggesting efficient drug target activity in all cell lines (Fig. 4A and Supplementary Fig. S7). Subsequently, immunofluorescence staining of TAK-981-treated SK-N-AS and KELLY cells showed an increased cytoplasmic fraction of SUMO1 and SUMO2/3 after 48 hours (Fig. 4B). Taken together, TAK-981 efficiently inhibits SUMOylation processes in NB cells.

**Figure 4.**
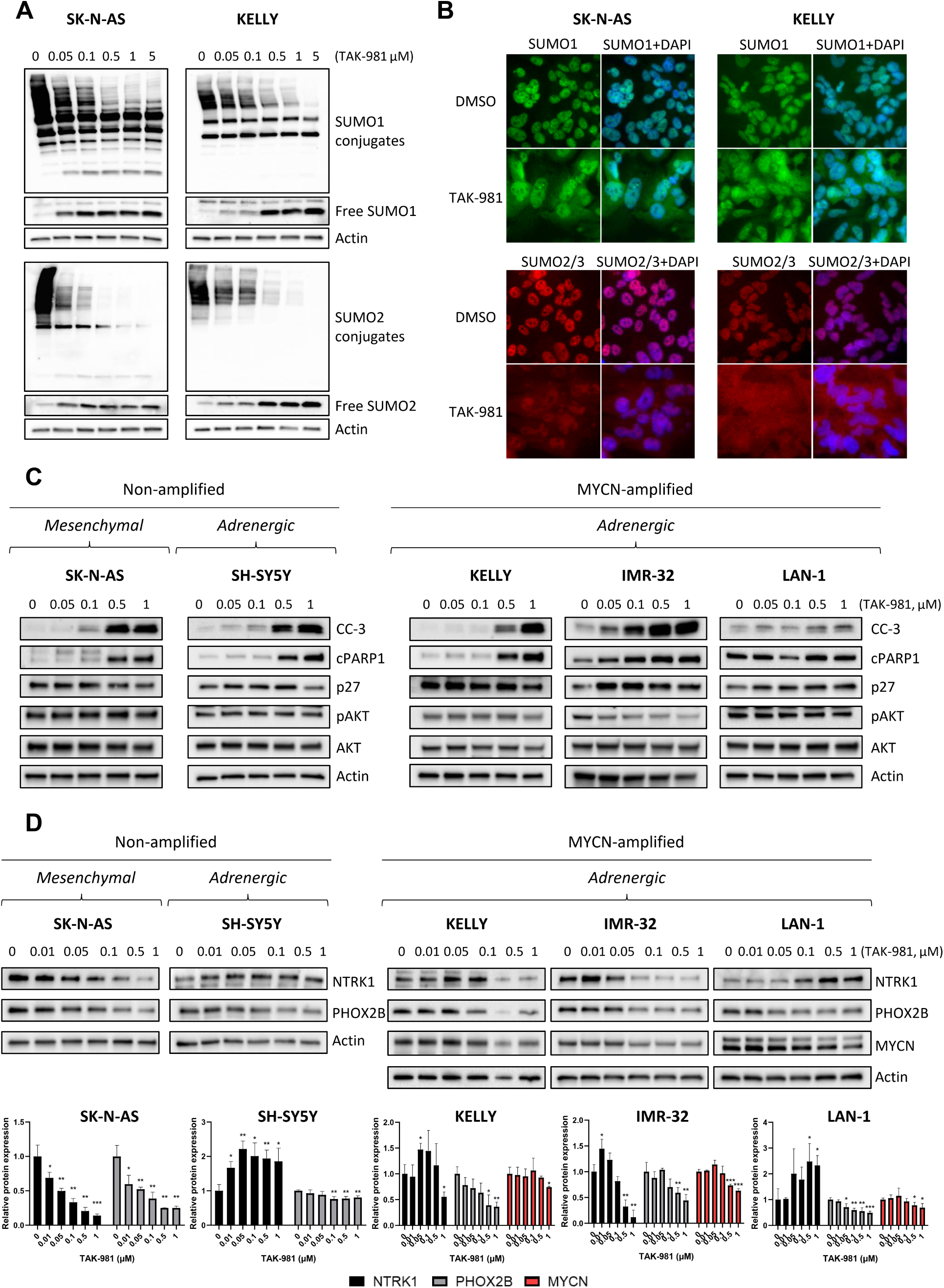
Inhibition of SUMOylation triggers apoptosis and influences differentiation pathways in NB cell lines. **(A)** Representative western blots of the conjugation levels of SUMO1 and SUMO2 in SK-N-AS and KELLY cells treated with increasing concentrations of TAK-981 for 48 hours. Longer exposures of SUMO1 and SUMO2 are shown below to illustrate the changes in the levels of free SUMO1 and SUMO2. **(B)** Immunofluorescence staining of SUMO1 and SUMO2/3 in SK-N-AS and KELLY cells treated with 0.5 µM TAK- 981 for 48 hours. DAPI was used for nuclear staining. **(C)** Representative western blots of cleaved caspase-3 (CC-3), cleaved PARP1 (cPARP), p27, pAKT, total AKT, and actin (loading control) levels in SK-N-AS, SH-SY5Y, KELLY, IMR-32 and LAN-1 cell lines treated with increasing concentrations of TAK-981 for 48 hours. **(D)** Representative western blots of the effect of TAK-981 on the levels of NTRK1 and PHOX2B in SK-N-AS, SH-SY5Y, KELLY, IMR-32, and LAN-1 cells. Additionally, MYCN level was evaluated from KELLY, IMR-32, and LAN-1. Bar graphs with SD depict pooled expression data (n=3) normalized to the loading control (actin) and further normalized to the vehicle control DMSO. Student’s t-test was used for statistical testing against the vehicle control (* p < 0.05, ** p < 0.01, *** p < 0.001).

Moreover, TAK-981 induced a concentration-dependent increase in cleaved caspase-3 and cleaved PARP levels in SK-N-AS, SH-SY5Y, KELLY, and IMR-32 cells after 48 hours (Fig. 4C), with IMR-32 being the most sensitive to the apoptotic effect of TAK-981, consistent with the 2D and 3D cell growth data. TAK-981 treatment resulted in a slight increase of cleaved caspase-3 expression also in the less SUMOylation inhibition sensitive LAN-1 cell line, whereas cleaved PARP levels remained unchanged. Moreover, the expression of the CDK/cell cycle inhibitor p27 was significantly upregulated in IMR-32 and LAN-1 cells, while its expression was unchanged in SK-N-AS, SH-SY5Y and KELLY cells. The phosphorylation level of AKT (s473), as an indicator of PI3K-AKT pro-survival pathway activity, was strongly downregulated in IMR-32 upon TAK-981 treatment, while these effects were not observed in the other cell lines after 48 hours. Overall, these data suggest that global SUMOylation inhibition induces apoptosis in both *MYCN*-amplified and non-amplified cells and may also promote cell cycle arrest in NB cells.

Next, we examined whether inhibition of SUMOylation affects differentiation-associated pathways *in vitro*. In line with patient mRNA co-expression data, TAK-981 increased the expression level of NTRK1 and reduced the expression of PHOX2B in SH-SY5Y and KELLY cells after 48 hours (Fig. 4D). Similarly, TAK-981 increased the expression of NTRK1 in the highly TAK-981-sensitive IMR-32 cells at a lower concentration; however, this induction of NTRK1 was not seen at the higher concentrations of TAK-981, likely due to decreased protein synthesis caused by extensive treatment-induced cell death. Expression levels of PHOX2B were reduced in a concentration-dependent manner in IMR-32 cells. In the more SUMO-inhibition-resistant LAN-1 cells, TAK-981 induced a dose-dependent upregulation of NTRK1 expression while simultaneously decreasing the levels of PHOX2B. Additionally, TAK-981 treatment moderately downregulated MYCN expression in all *MYCN*- amplified cell lines. In contrast, TAK-981 treatment resulted in downregulation of both NTRK1 and PHOX2B expression in SK-N-AS cells, possibly due to their mesenchymal phenotype. Taken together, these data suggest that inhibition of SUMOylation may promote partial differentiation in NB cell lines with an adrenergic phenotype.

### ATRA downregulates the expression of SUMOylation-promoting factors and potentiates the effect of TAK-981

Next, we investigated whether the differentiation-inducing agent ATRA affects the expression levels of SUMOylation-promoting factors in NB cells. Analysis of previously published RNA-seq data(*35*) showed decreased levels of *SAE1*, *SAE2*, and *UBE2I* in *MYCN*-amplified SK-N-BE(2)C and NGP cells, as well as in non-amplified NBL-S cells after 6 days of ATRA treatment (Fig. 5A). *UBE2I* expression decreased similarly across all cell lines, whereas *SAE1* and *SAE2* downregulation was significantly more prominent in ATRA-responsive SK-N-BE(2)C and NGP cells compared to ATRA-resistant NBL-S cells. ATRA also reduced expression of *SAE1*, *SAE2*, and *UBE2I* in SH-SY5Y-E and SK-N-SH cells(*36*) (Supplementary Fig. S8A). Moreover, differential gene expression analysis of an independent dataset(*37*) confirmed downregulation of SUMOylation-promoting factors and negative enrichment of the REACTOME_SUMOYLATION gene set in ATRA-treated CHP-134 cells after 6 days (Fig. 5B). Consistently, ATRA reduced SAE1, SAE2, and UBC9 protein levels after 6 days in the ATRA-responsive KELLY cell line (Fig. 5C). Overall, these data indicate that the growth-inhibitory effects of ATRA are accompanied by downregulation of SUMOylation-promoting factors.

**Figure 5.**
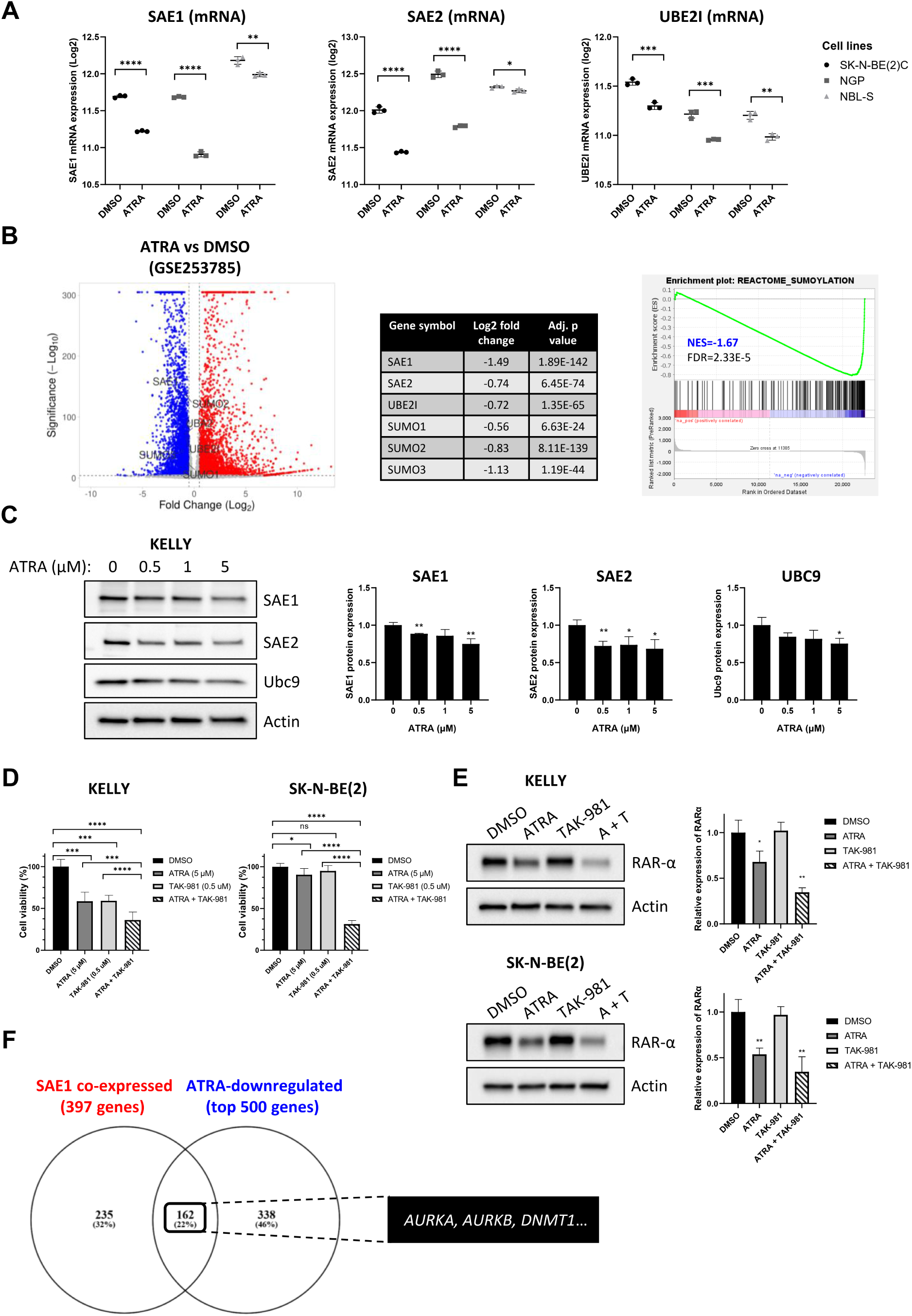
ATRA downregulates the expression of SUMOylation-promoting factors and potentiates the effect of TAK-981 in NB cell lines. **(A)** Scatter dot plots with SD depicting the expression (log2) of *SAE1*, *SAE2*, and *UBE2I* between DMSO and 5 µM ATRA treatment groups after 6 days in SK-N-BE(2)C (black; n=3), NGP (grey; n=3), and NBL-S (light grey; n=3) cells. Analysis was done from GSE155000. Statistical significance was determined with Student’s t-test (* p < 0.05, ** p < 0.01, *** p < 0.001, **** p < 0.0001). **(B)** Volcano plot of differentially expressed genes (adjusted p value <0.001 and log2 fold change >0.5) after 6 days of ATRA treatment in CHP-134 cells (GSE253785) and a table showing log2 fold changes and adjusted p values of SUMOylation-promoting factors. GSEA plot displays the NES and FDR for the gene set REACTOME_SUMOYLATION in ATRA-treated cells. **(C)** Representative western blots of the effect of ATRA (0–5 µM) on the levels of SAE1, SAE2, and UBC9 in KELLY cells after 6 days of treatment. Bar graphs with SD depict pooled western blot densitometry data (n=3) normalized to the loading control (actin) and further normalized to vehicle control DMSO. Student’s t-test was used for statistical testing (* p < 0.05, ** p < 0.01). **(D)** Bar graphs with SD depicting the cell viability of SK-N-BE(2) and KELLY cells treated with ATRA (5 µM), TAK-981 (0.5 µM) and the combination of ATRA and TAK-981 after 4 days. Viability of cells was determined with the MTS assay. **(E)** Representative western blots of the effect of ATRA (5 µM), TAK-981 (0.5 µM), and the combination of ATRA and TAK- 981 on the expression level of RARα in SK-N-BE(2) and KELLY cells after 4 days. Bar graphs with SD depict pooled western blot densitometry data (n=3) normalized to loading control (actin) and further normalized to vehicle control DMSO. Student’s t-test was used for statistical testing (* p < 0.05, ** p < 0.01). **(F)** Venn diagram showing the overlap of the top 500 genes downregulated by ATRA in CHP-134 cells (analyzed from GSE253785) and the list of the 397 commonly co-expressed genes with *SAE1* identified from the top 1000 positively co-expressed genes in Kocak, SEQC, Cangelosi, and Versteeg NB patient cohorts (Supplementary table S3).

Next, we studied the combined effects of ATRA and TAK-981 in KELLY and SK-N-BE(2) cells. Interestingly, the cell viability in both cell lines was most significantly decreased after 4 days following combination treatment compared with either agent alone (Fig. 5D, Supplementary Fig. S8B). At the molecular level, the protein expression of retinoic acid receptor alpha (RARα) is known to decrease upon ATRA-induced activation of retinoic acid signaling(*43*). The strongest downregulation of RARα protein expression was observed in the ATRA - TAK-981 -combination treatment after 4 days, indicating for potential enhanced pathway activation (Fig. 5E). Taken together, these findings suggest that inhibition of SUMOylation may potentiate the growth-inhibiting and differentiation-inducing effects of retinoic acids in NB.

Next, we compared the genes that were downregulated by ATRA in CHP-134 cells (top 500 genes) to genes positively co-expressed with *SAE1* in NB patient cohorts (397 genes, see Supplementary table S3) and observed a significant overlap (162 genes, 22%, Supplementary table S4) in the gene lists (Fig. 5F). Not surprisingly, genes such as the cycling neuroblast markers *MKI67* and *TOP2A* were found in the list further supporting interaction of retinoic acid signaling and SUMOylation pathway in regulation of NB cell proliferation states. To identify additional genes and pathways that could be targeted synergistically with SUMOylation inhibition we chose select promising genes for further study, including aurora kinase A (*AURKA*) and aurora kinase B (*AURKB*), given their established role in NB pathogenesis(*13*, *44*), as well as DNA methyltransferase 1 (*DNMT1*), which has been reported to be positively regulated by SUMOylation(*45*, *46*).

### *SAE1* expression associates with high levels of DNMT1 and TAK-981 synergizes with DNMT inhibition in NB

To identify combination partners for TAK-981, we first tested whether TAK-981 synergizes with aurora kinase inhibition in 3D-cultured *MYCN*-amplified IMR-32 cells using the CellTiter-Glo assay for cell viability readout. The synergy was determined using SynergyFinder, which calculates the synergy score based on drug sensitivity and resistance score (DSS) values of the drugs; when the score is larger than 10 the interaction between two drugs is likely to be synergistic(*38*). TAK-981 showed no synergy with either the pan-aurora kinase inhibitor tozasertib or the aurora A-selective inhibitor alisertib (Synergy scores 1.295 for tozasertib combination and -0.457 for alisertib, Supplementary Fig. S9A).

Therefore, we focused on DNMT1, which showed strong co-expression with *SAE1* at the mRNA level in NB patient samples (Fig. 6A) and at the protein level in NB cell lines (Fig. 6B). Combination treatment with TAK-981 and the DNMT inhibitor decitabine (5-Aza-2’- deoxycytidine) resulted in a marked synergy (score 14.974) in IMR-32 3D spheroids (Fig. 6C, Supplementary Fig. S9B). Mechanistically, this combination resulted in accumulation of DNA damage and induction of apoptosis, as indicated by increased γH2AX and cleaved PARP levels in the western blot (Fig. 6D). Similar effects were seen in *MYCN*-non-amplified SH-SY5Y cells, showing that the synergistic interaction between SUMOylation inhibition and DNMT inhibition is not limited to *MYCN*-amplified NB.

**Figure 6.**
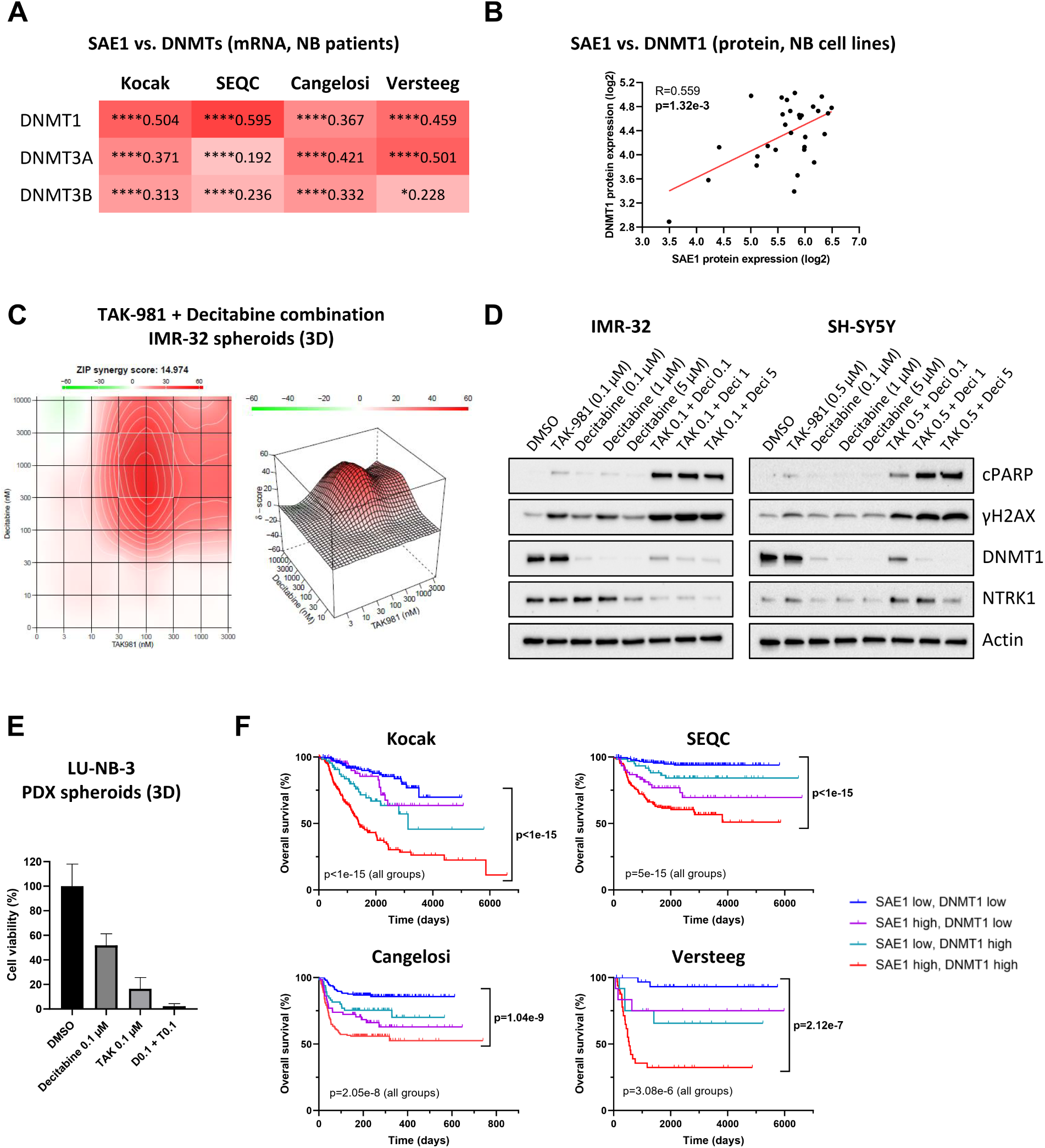
Inhibition of SUMOylation synergizes with the DNMT inhibitor decitabine. **(A)** Co-expression of *SAE1* with *DNMT1*, *DNMT3A*, and *DNMT3B* in Kocak, SEQC, Cangelosi, and Versteeg cohorts determined with Pearson correlation analysis (* p < 0.05, **** p < 0.0001). **(B)** Correlation of SAE1 and DNMT1 protein levels in human NB cell lines of the ProCan-DepMapSanger dataset(*69*) determined with Pearson correlation analysis. **(C)** IMR-32 cells were grown in 3D and treated with concentration series of TAK-981 and DNMT inhibitor decitabine to determine the ZIP synergy score based on cell viability after 72 hours of treatment. **(D)** Representative western blots of the effect of TAK-981 (0.1 µM in IMR-32; 0.5 µM in SH-SY5Y) and decitabine (0.1, 1, and 5 µM) combination treatment on the levels of cPARP, γH2AX, DNMT1, and NTRK1 after 48 hours. Actin was used as the loading control. **(E)** Representative bar graph of the effect of TAK-981 (0.1 µM), decitabine (0.1 µM), and their combination on the cell viability of 3D-cultured LU-NB-3 PDX spheroids after 72 hours measured with the WST-8 assay. **(F)** Kaplan-Meier plots of the combined expression of *SAE1* and *DNMT1* on OS of NB patients in the Kocak, SEQC, Cangelosi, and Versteeg cohorts. Median mRNA expression value was used as a cutoff for group stratification and the patients were divided to four subgroups. Statistical significance was determined with the log-rank test.

To evaluate this combination in a more physiologically relevant setting, we utilized *MYCN*-amplified LU-NB-3 cells derived from a PDX NB model(*47*, *48*). While TAK-981 monotherapy significantly inhibited growth of LU-NB-3 cells grown in 3D culture, the combination with decitabine further enhanced this effect, supporting synergistic activity also in PDX-derived NB cells (Fig. 6E).

Consistent with these findings, NB patients expressing high levels of both *SAE1* and *DNMT1* had significantly poorer survival (Kocak p=<1e-15, SEQC p=1e-15, Cangelosi p=1.04e-9, Versteeg p=2.12e-7), whereas patients with low expression of both genes formed the subgroup with the most favorable prognosis across all analyzed cohorts, further supporting the rationale for co-targeting of SUMOylation and DNA methylation in NB (Fig. 6F).

## Discussion

Although some NBs are indolent, aggressive high-risk NB has poor prognosis despite intensive treatment. Improved prognostic markers and more efficient therapies are needed. Here we have extensively characterized the prognostic power and the therapeutic potential of targeting the SUMO pathway in NB by using multiple patient datasets representing different patient cohorts and by performing experiments with a panel of NB cell lines. We show that increased expression of SUMOylation-promoting SUMO E1 and E2 enzymes *SAE1*/*2* and *UBE2I*, as well as deSUMOylase *SENP2*, and the decreased expression of deSUMOylases *SENP6* and *SENP7* associate with poor prognosis and aggressive features of NB. Accordingly, inhibition of global SUMOylation with SAE inhibitor TAK-981 reduces growth of NB cells by inducing cell cycle arrest, apoptosis, and the regulation of differentiation-associated pathways. Moreover, high levels of *SAE1*/*2* and *UBE2I* also associate with the downregulation of late differentiation markers of NB. Interestingly, low expression of late neuroblast differentiation markers in NB cell lines correlates with increased sensitivity to SUMOylation inhibition, and TAK-981 potentiates the growth-inhibitory effects of retinoic acid in NB cells. In addition, we demonstrate synergistic activity of TAK-981 and DNMT inhibitor decitabine in NB cell lines and proof-of-concept evidence using a NB PDX model.

Due to the unmet need for better therapies in NB, search after druggable driver mutations has been active, but the breakthroughs entering routine clinical use have been slow. The relatively low mutation burden in NB has also hampered the expectations for the efficacy of immune checkpoint inhibitors(*49*). Ongoing registered trials are testing, for example, drugs targeting cell surface disialogangliosides, different kinases and cell survival pathways, histone deacetylases, and the protein degradation machinery. Moreover, many selective targeted therapies guided by genetic tests have been evaluated in the multicenter pediatric MATCH trial (NCT03155620). Recent results with ALK-targeting drug lorlatinib are promising and support its transition to phase III trials(*16*). Intriguingly, our data suggests that inhibition of global SUMOylation may have therapeutic potential in NB as a novel approach that may also potentiate the effect of differentiation-inducing retinoic acids that are in clinical use. The first-in-class drug TAK-981, which inhibits the SUMO E1-activating enzyme, has been shown to be tolerable in adult patients in early phase trials, although adverse effects such as fatigue, nausea, headache, pyrexia, and cytokine syndrome have been reported(*50*). The possible side effects of SUMOylation inhibition to the developing tissues in children are not known and should be carefully considered prior any translation towards clinics.

Our data suggest that both cell survival and differentiation pathways in NB are regulated by SUMOylation. The mechanism of action for inhibitors that reduce global SUMOylation in cancer is not fully understood, but they likely act by interfering with SUMOylation of substrates important for transformed phenotype and by disrupting cellular processes tightly regulated by SUMO, such as transcription, DNA repair, cell cycle progression, and cell signaling(*17*, *19*). Intriguingly, inhibition of SUMOylation by TAK-981 has also been suggested to be a novel way to activate the immune response to fight cancer cells(*51*). Little has been reported about SUMO in NB, but a few preclinical studies have suggested roles for SUMOylation in regulating the functions of specific substrates such as the deubiquitinating enzyme CYLD and Ephrin receptor B1(*52*, *53*), and the deSUMOylase SENP1 has been suggested to regulate migration of NB cells(*54*). Moreover, a study conducted in three NB patients with stage 4 disease and deleted 11q identified a gained region in ch17q comprising *SUMO2*, *BIRC5*, *BRCA1*, *PRKCA*, and *GPS1*, suggesting copy number gains of SUMO2 in high-risk NB(*55*). Although not specifically addressed in NB, MYCN has also been connected to SUMOylation(*23*). *In vitro* experiments with NB cells show that SUMOylation is stress-regulated and important for cell division as well as for cell survival in low oxygen and glucose conditions, supporting our findings(*56–58*). The general role of the SUMO pathway and its components, as well as SUMO substrates implicated in oncogenesis, needs to be explored in more detail in NB.

Although c-MYC and MYCN have some functional differences, we expected that *MYCN* amplification status would predict sensitivity to TAK-981 in NB based on previous data showing that aberrant upregulation of SUMOylation is essential for the pathogenesis of some c-MYC-dependent cancers(*24–26*). We also found that expression levels of SUMOylation- promoting enzymes increase in *MYCN*-amplified NB patients and mouse models. However, our data suggests that in the context of NB, the *MYCN*-amplified cells were not significantly more sensitive to inhibition of SUMOylation than the *MYCN*-non-amplified cells, suggesting that *MYCN* amplification may not be a general tumor-agnostic predictive marker of sensitivity to inhibition of SUMOylation. After treatment with TAK-981, MYCN levels were slightly reduced in all tested cell lines, but this reduction did not correlate with treatment response. In our panel of NB cells, levels of SAE1/2, UBC9, or SUMO proteins also did not seem to correlate with sensitivity to TAK-981. Interestingly, *SUMO2* and *PC2* amplifications were detected in IMR-32 and SK-N-AS cell lines, both of which were relatively sensitive to TAK- 981. *SUMO2* and *PC2* amplifications were also found in the small cohort of the TARGET study in 8.5% and 10.2% of patients, respectively, and their prevalence in other larger cohorts should be assessed. We derived a late neuroblast signature score, which significantly correlated with cell line sensitivity to TAK-981 inhibition. The future goal is to further elucidate the predictive markers that determine sensitivity to global SUMOylation inhibitors in NB.

NBs are thought to arise from neural crest-derived sympathetic progenitor cells, not fully committing to differentiation programs forming sympathetic neuronal cells and chromaffin neuroendocrine cells, but instead prematurely exiting at different phases of lineage-specific differentiation path(*40*). Our data show that TAK-981 treatment alters NTRK1 and PHOX2B expression and link high expression of SUMOylation-promoting enzymes to the undifferentiated proliferative subpopulation of NB cells in patients, suggesting that the SUMO machinery is likely involved in regulation of adrenergic gene expression programs. Numerous studies have previously linked SUMOylation to cell fate decisions and cellular differentiation via e.g. regulatory constraints it exerts over transcription factors and chromatin remodeling complexes(*59*). Regulation of protein SUMOylation is highly dynamic during differentiation, and SUMO paralogs may have distinct and opposing roles(*60–62*). In NB, retinoic acid is known to induce differentiation, and its derivatives are used in treatments(*1*). Retinoic acid is suggested to alter the activity of transcription factors regulating adrenergic gene expression programs, thereby promoting differentiation of neuroblasts into mature sympathetic neuronal cells(*35*). Retinoic acid binds to RARs, nuclear hormone receptors regulating gene transcription after binding to retinoic acid response elements in the promoter regions of target genes. RARα/RARγ double knockout mice exhibit neural crest defects during early embryogenesis, suggesting that RARs function as regulators of cells arising from neural crest(*63*, *64*). SUMOylation has previously been suggested in the regulation of RAR transcriptional activity, providing one possible mechanism behind the effects of TAK-981 in NB(*65*). We observed that co-treatment with ATRA and TAK-981 resulted in downregulation of RARα expression and strong growth reduction in NB cells, suggesting that TAK-981 may potentiate the effects of ATRA. Interestingly, both ATRA(*66*, *67*) and SUMOylation inhibition(*45*, *46*) have been shown to inhibit the expression or activity of DNMTs, and we show that combining SUMOylation inhibition with the DNMT inhibitor decitabine also results in synergistic reduction in NB cell viability via accumulation of DNA damage.

Our data reveals that high levels of the deSUMOylates *SENP6* and *SENP7* associate with improved survival in NB, suggesting potential tumor suppressive functions. Neither of these deSUMOylases nor their substrates have been characterized in NB, but previous data in other contexts also supports their anti-tumorigenic roles(*17*). Interestingly, SENP7 has previously been linked to regulation of neuronal differentiation(*68*), which could be related to its favorable prognostic impact in NB.

In conclusion, we elucidate to our knowledge for the first time the prognostic significance of SUMO pathway components and promising therapeutic potential in the inhibition of SUMOylation in NB, warranting further studies. Future goals include the characterization of key SUMO substrates in NB, development of assays to detect the activation status of the SUMO pathway in tissues, testing the activity of inhibition of global SUMOylation in combination therapies *ex vivo,* and identifying biomarkers that predict response to SUMO inhibition.

## Data Availability

Data are available from authors upon reasonable request.

## Acknowledgements

We thank Medical Bioinformatics Centre (Turku Bioscience Centre and supported by Biocenter Finland) for consultations. We thank Minna Santanen and Barbara Vähätalo-Ramos for excellent technical assistance. We thank Dr. Baraa Abuasaker and Milla Hollmén for valuable comments and discussions. FIMM High Throughput Biomedicine core unit (University of Helsinki, Finland; Biocenter Finland and EU OpenScreen), and especially Dr. Laura Turunen and Dr. Swapnil Potdar are acknowledged for their expertise in drug plate preparations and drug synergy testing. We thank Zebrafish Core (Turku Bioscience and supported by Biocenter Finland) and Sk Hasan for technical assistance with the *in vivo* zebrafish experiment.

## Author contributions

AK and MS conceptualized the study. AK, VV, and MS designed the *in silico* analyses. AK, VV, and LKB carried out *in silico* analyses. AK, SS, and MS designed the *in vitro* experiments. AK, SS, and LKB carried out the *in vitro* experiments. SK and VP designed, carried out, and analyzed the drug synergy experiments. IP designed and supervised the *in vivo* zebrafish experiment. AK analyzed the *in vivo* data. DB provided materials and help with the patient- derived organoid experiments. AK and MS wrote the manuscript. MS acquired the funding and supervised the study. All authors have read, commented, and approved the final version of the manuscript.

## References

1. Maris JM, Recent Advances in Neuroblastoma. N. Engl. J. Med. 362, 2202–2211 (2010).

2. H. Shimada, I. M. Ambros, L. P. Dehner, J. Hata, V. V. Joshi, B. Roald, D. O. Stram, R. B. Gerbing, J. N. Lukens, K. K. Matthay, R. P. Castleberry, The International Neuroblastoma Pathology Classification (the Shimada system). Cancer 86, 364–372 (1999).

3. M. S. Irwin, A. Naranjo, F. F. Zhang, S. L. Cohn, W. B. London, J. M. Gastier-Foster, N. C. Ramirez, R. Pfau, S. Reshmi, E. Wagner, J. Nuchtern, S. Asgharzadeh, H. Shimada, J. M. Maris, R. Bagatell, J. R. Park, M. D. Hogarty, Revised Neuroblastoma Risk Classification System: A Report From the Children’s Oncology Group. J. Clin. Oncol. 39, 3229–3241 (2021).

4. B. Qiu, K. K. Matthay, Advancing therapy for neuroblastoma. Nature Research (2022). 10.1038/s41571-022-00643-z.

5. D. Trochet, F. Bourdeaut, I. Janoueix-Lerosey, A. Deville, L. De Pontual, G. Schleiermacher, C. Coze, N. Philip, T. Frébourg, A. Munnich, S. Lyonnet, O. Delattre, J. Amiel, Germline Mutations of the Paired-Like Homeobox 2B (PHOX2B) Gene in Neuroblastoma. Am. J. Hum. Genet. 74, 761–764 (2004).

6. Y. P. Mosse, M. Laudenslager, D. Khazi, A. J. Carlisle, C. L. Winter, E. Rappaport, J. M. Maris, Germline PHOX2B mutation in hereditary neuroblastoma. University of Chicago Press (2004). 10.1086/424530.

7. I. Janoueix-Lerosey, D. Lequin, L. Brugières, A. Ribeiro, L. De Pontual, V. Combaret, V. Raynal, A. Puisieux, G. Schleiermacher, G. Pierron, D. Valteau-Couanet, T. Frebourg, J. Michon, S. Lyonnet, J. Amiel, O. Delattre, Somatic and germline activating mutations of the ALK kinase receptor in neuroblastoma. Nature 455, 967–970 (2008).

8. Y. P. Mossé, M. Laudenslager, L. Longo, K. A. Cole, A. Wood, E. F. Attiyeh, M. J. Laquaglia, R. Sennett, J. E. Lynch, P. Perri, G. Laureys, F. Speleman, C. Kim, C. Hou, H. Hakonarson, A. Torkamani, N. J. Schork, G. M. Brodeur, G. P. Tonini, E. Rappaport, M. Devoto, J. M. Maris, Identification of ALK as a major familial neuroblastoma predisposition gene. Nature 455, 930–935 (2008).

9. G. M. Brodeur, R. C. Seeger, M. Schwab, H. E. Varmus, J. Michael Bishop, Amplification of N-myc in untreated human neuroblastomas correlates with advanced disease stage. Science 224, 1121–1124 (1984).

10. N. K. V. Cheung, J. Zhang, C. Lu, M. Parker, A. Bahrami, S. K. Tickoo, A. Heguy, A. S. Pappo, S. Federico, J. Dalton, I. Y. Cheung, L. Ding, R. Fulton, J. Wang, X. Chen, J. Becksfort, J. Wu, C. A. Billups, D. Ellison, E. R. Mardis, R. K. Wilson, J. R. Downing, M. A. Dyer, Association of age at diagnosis and genetic mutations in patients with neuroblastoma. JAMA 307, 1062–1071 (2012).

11. T. J. Pugh, O. Morozova, E. F. Attiyeh, S. Asgharzadeh, J. S. Wei, D. Auclair, S. L. Carter, K. Cibulskis, M. Hanna, A. Kiezun, J. Kim, M. S. Lawrence, L. Lichenstein, A. Mckenna, C. S. Pedamallu, A. H. Ramos, E. Shefler, A. Sivachenko, C. Sougnez, C. Stewart, A. Ally, I. Birol, R. Chiu, R. D. Corbett, M. Hirst, S. D. Jackman, B. Kamoh, A. H. Khodabakshi, M. Krzywinski, A. Lo, R. A. Moore, K. L. Mungall, J. Qian, A. Tam, N. Thiessen, Y. Zhao, K. A. Cole, M. Diamond, S. J. Diskin, Y. P. Mosse, A. C. Wood, L. Ji, R. Sposto, T. Badgett, W. B. London, Y. Moyer, J. M. Gastier-Foster, M. A. Smith, J. M. G. Auvil, D. S. Gerhard, M. D. Hogarty, S. J. M. Jones, E. S. Lander, S. B. Gabriel, G. Getz, R. C. Seeger, J. Khan, M. A. Marra, M. Meyerson, J. M. Maris, The genetic landscape of high-risk neuroblastoma. Nat. Genet. 45, 279–284 (2013).

12. C. Rosswog, J. Fassunke, A. Ernst, B. Schömig-Markiefka, S. Merkelbach-Bruse, C. Bartenhagen, M. Cartolano, S. Ackermann, J. Theissen, M. Blattner-Johnson, B. Jones, K. Schramm, J. Altmüller, P. Nürnberg, M. Ortmann, F. Berthold, M. Peifer, R. Büttner, F. Westermann, J. H. Schulte, T. Simon, B. Hero, M. Fischer, Genomic ALK alterations in primary and relapsed neuroblastoma. Br. J. Cancer 128, 1559–1571 (2023).

13. T. Otto, S. Horn, M. Brockmann, U. Eilers, L. Schüttrumpf, N. Popov, A. M. Kenney, J. H. Schulte, R. Beijersbergen, H. Christiansen, B. Berwanger, M. Eilers, Stabilization of N-Myc Is a Critical Function of Aurora A in Human Neuroblastoma. Cancer Cell 15, 67–78 (2009).

14. E. Chipumuro, E. Marco, C. L. Christensen, N. Kwiatkowski, T. Zhang, C. M. Hatheway, B. J. Abraham, B. Sharma, C. Yeung, A. Altabef, A. Perez-Atayde, K. K. Wong, G. C. Yuan, N. S. Gray, R. A. Young, R. E. George, CDK7 inhibition suppresses super-enhancer-linked oncogenic transcription in MYCN-driven cancer. Cell 159, 1126–1139 (2014).

15. A. K. Brenner, M. W. Gunnes, Therapeutic targeting of the anaplastic lymphoma kinase (ALK) in neuroblastoma—A comprehensive update. MDPI (2021). 10.3390/pharmaceutics13091427.

16. K. C. Goldsmith, J. R. Park, K. Kayser, J. Malvar, Y. Y. Chi, S. G. Groshen, J. G. Villablanca, K. Krytska, L. M. Lai, P. T. Acharya, F. Goodarzian, B. Pawel, H. Shimada, S. Ghazarian, L. States, L. Marshall, L. Chesler, M. Granger, A. V. Desai, R. Mody, D. A. Morgenstern, S. Shusterman, M. E. Macy, N. Pinto, G. Schleiermacher, K. Vo, H. C. Thurm, J. Chen, M. Liyanage, G. Peltz, K. K. Matthay, E. R. Berko, J. M. Maris, A. Marachelian, Y. P. Mossé, Lorlatinib with or without chemotherapy in ALK-driven refractory/relapsed neuroblastoma: phase 1 trial results. Nat. Med. 29, 1092–1102 (2023).

17. A. Kukkula, V. K. Ojala, L. M. Mendez, L. Sistonen, K. Elenius, M. Sundvall, Therapeutic Potential of Targeting the SUMO Pathway in Cancer. MDPI (2021). 10.3390/cancers13174402.

18. J. S. Kroonen, A. C. O. Vertegaal, Targeting SUMO Signaling to Wrestle Cancer. Cell Press (2021). 10.1016/j.trecan.2020.11.009.

19. R. T. Hay, SUMO: A history of modification. Cell Press (2005). 10.1016/j.molcel.2005.03.012.

20. A. Pichler, C. Fatouros, H. Lee, N. Eisenhardt, SUMO conjugation - A mechanistic view. Walter de Gruyter GmbH (2017). 10.1515/bmc-2016-0030.

21. M. Sundvall, Role of ubiquitin and SUMO in intracellular trafficking. Curr. Issues Mol. Biol. 35, 99–108 (2020).

22. M. Kalkat, P. K. Chan, A. R. Wasylishen, T. Srikumar, S. S. Kim, R. Ponzielli, D. P. Bazett- Jones, B. Raught, L. Z. Penn, Identification of c-MYC SUMOylation by mass spectrometry. PLoS One 9 (2014).

23. A. Sabò, M. Doni, B. Amati, SUMOylation of Myc-family proteins. PLoS One 9 (2014).

24. J. D. Kessler, K. T. Kahle, T. Sun, K. L. Meerbrey, M. R. Schlabach, E. M. Schmitt, S. O. Skinner, Q. Xu, M. Z. Li, Z. C. Hartman, M. Rao, P. Yu, R. Dominguez-Vidana, A. C. Liang, N. L. Solimini, R. J. Bernardi, B. Yu, T. Hsu, I. Golding, J. Luo, C. K. Osborne, C. J. Creighton, S. G. Hilsenbeck, R. Schiff, C. A. Shaw, S. J. Elledge, T. F. Westbrook, A SUMOylation-dependent transcriptional subprogram is required for Myc-driven tumorigenesis. Science 335, 348–353 (2012).

25. A. Hoellein, M. Fallahi, S. Schoeffmann, S. Steidle, F. X. Schaub, M. Rudelius, I. Laitinen, L. Nilsson, A. Goga, C. Peschel, J. A. Nilsson, J. L. Cleveland, U. Keller, Myc-induced SUMOylation is a therapeutic vulnerability for B-cell lymphoma. Blood 124, 2081–2090 (2014).

26. A. Rabellino, M. Melegari, V. S. Tompkins, W. Chen, B. G. Van Ness, J. Teruya-Feldstein, M. Conacci-Sorrell, S. Janz, P. P. Scaglioni, PIAS1 Promotes Lymphomagenesis through MYC Upregulation. Cell Rep. 15, 2266–2278 (2016).

27. A. Biederstädt, Z. Hassan, C. Schneeweis, M. Schick, L. Schneider, A. Muckenhuber, Y. Hong, G. Siegers, L. Nilsson, M. Wirth, Z. Dantes, K. Steiger, K. Schunck, S. Langston, H. P. Lenhof, A. Coluccio, F. Orben, J. Slawska, A. Scherger, D. Saur, S. Müller, R. Rad, W. Weichert, J. Nilsson, M. Reichert, G. Schneider, U. Keller, SUMO pathway inhibition targets an aggressive pancreatic cancer subtype. Gut 69, 1472–1482 (2020).

28. S. P. Langston, S. Grossman, D. England, R. Afroze, N. Bence, D. Bowman, N. Bump, R. Chau, B. C. Chuang, C. Claiborne, L. Cohen, K. Connolly, M. Duffey, N. Durvasula, S. Freeze, M. Gallery, K. Galvin, J. Gaulin, R. Gershman, P. Greenspan, J. Grieves, J. Guo, N. Gulavita, S. Hailu, X. He, K. Hoar, Y. Hu, Z. Hu, M. Ito, M. S. Kim, S. W. Lane, D. Lok, A. Lublinsky, W. Mallender, C. McIntyre, J. Minissale, H. Mizutani, M. Mizutani, N. Molchinova, K. Ono, A. Patil, M. Qian, J. Riceberg, V. Shindi, M. D. Sintchak, K. Song, T. Soucy, Y. Wang, H. Xu, X. Yang, A. Zawadzka, J. Zhang, S. M. Pulukuri, Discovery of TAK- 981, a First-in-Class Inhibitor of SUMO-Activating Enzyme for the Treatment of Cancer. J. Med. Chem., doi: 10.1021/acs.jmedchem.0c01491 (2021).

29. H. Kocak, S. Ackermann, B. Hero, Y. Kahlert, A. Oberthuer, D. Juraeva, F. Roels, J. Theissen, F. Westermann, H. Deubzer, V. Ehemann, B. Brors, M. Odenthal, F. Berthold, M. Fischer, Hox-C9 activates the intrinsic pathway of apoptosis and is associated with spontaneous regression in neuroblastoma. Cell Death Dis. 4, e586–e586 (2013).

30. Z. Su, H. Fang, H. Hong, L. Shi, W. Zhang, W. Zhang, Y. Zhang, Z. Dong, L. J. Lancashire, M. Bessarabova, X. Yang, B. Ning, B. Gong, J. Meehan, J. Xu, W. Ge, R. Perkins, M. Fischer, W. Tong, An investigation of biomarkers derived from legacy microarray data for their utility in the RNA-seq era. Genome Biol. 15, 523 (2014).

31. J. J. Molenaar, J. Koster, D. A. Zwijnenburg, P. Van Sluis, L. J. Valentijn, I. Van Der Ploeg, M. Hamdi, J. Van Nes, B. A. Westerman, J. Van Arkel, M. E. Ebus, F. Haneveld, A. Lakeman, L. Schild, P. Molenaar, P. Stroeken, M. M. Van Noesel, I. Øra, E. E. Santo, H. N. Caron, E. M. Westerhout, R. Versteeg, Sequencing of neuroblastoma identifies chromothripsis and defects in neuritogenesis genes. Nature 483, 589–593 (2012).

32. D. Cangelosi, M. Morini, N. Zanardi, A. R. Sementa, M. Muselli, M. Conte, A. Garaventa, U. Pfeffer, M. C. Bosco, L. Varesio, A. Eva, Hypoxia predicts poor prognosis in neuroblastoma patients and associates with biological mechanisms involved in telomerase activation and tumor microenvironment reprogramming. Cancers (Basel). 12, 1–45 (2020).

33. N. Bonine, V. Zanzani, A. Van Hemelryk, B. Vanneste, C. Zwicker, T. Thoné, S. Roelandt, S. L. Bekaert, J. Koster, I. Janoueix-Lerosey, C. Thirant, S. Van Haver, S. S. Roberts, L. M. Mus, B. De Wilde, N. Van Roy, C. Everaert, F. Speleman, V. Vermeirssen, C. L. Scott, K. De Preter, NBAtlas: A harmonized single-cell transcriptomic reference atlas of human neuroblastoma tumors. Cell Rep. 43, 114804 (2024).

34. L. C. Heukamp, T. Thor, A. Schramm, K. De Preter, C. Kumps, B. De Wilde, A. Odersky, M. Peifer, S. Lindner, A. Spruessel, F. Pattyn, P. Mestdagh, B. Menten, S. Kuhfittig-Kulle, A. Künkele, K. König, L. Meder, S. Chatterjee, R. T. Ullrich, S. Schulte, J. Vandesompele, F. Speleman, R. Büttner, A. Eggert, J. H. Schulte, Targeted expression of mutated ALK induces neuroblastoma in transgenic mice. Sci. Transl. Med. 4 (2012).

35. M. W. Zimmerman, A. D. Durbin, S. He, F. Oppel, H. Shi, T. Tao, Z. Li, A. Berezovskaya, Y. Liu, J. Zhang, R. A. Young, B. J. Abraham, A. T. Look, Retinoic acid rewires the adrenergic core regulatory circuitry of childhood neuroblastoma. Sci. Adv. 7 (2021).

36. Y. Nishida, N. Adati, R. Ozawa, A. Maeda, Y. Sakaki, T. Takeda, Identification and classification of genes regulated by phosphatidylinositol 3-kinase- and TRKB-mediated signalling pathways during neuronal differentiation in two subtypes of the human neuroblastoma cell line SH-SY5Y. BMC Res. Notes 1, 95 (2008).

37. M. Pan, Y. Zhang, W. C. Wright, X. Liu, B. Passaia, D. Currier, J. Low, R. H. Chapple, J. A. Steele, J. P. Connelly, B. Ju, E. Plyler, M. Lu, A. J. Loughran, L. Yang, B. J. Abraham, S. M. Pruett-Miller, B. Freeman, G. E. Campbell, M. A. Dyer, T. Chen, E. Stewart, S. Koo, H. Sheppard, J. Easton, P. Geeleher, Bone morphogenetic protein (BMP) signaling determines neuroblastoma cell fate and sensitivity to retinoic acid. Nat. Commun. 16, 2036 (2025).

38. A. Ianevski, A. K. Giri, T. Aittokallio, SynergyFinder 2.0: Visual analytics of multi-drug combination synergies. Nucleic Acids Res. 48, W488–W493 (2021).

39. V. Virtanen, K. Paunu, A. Kukkula, S. Niva, Y. Junila, M. Toriseva, T. Jokilehto, S. Mäkelä, R. Huhtaniemi, M. Poutanen, I. Paatero, M. Sundvall, Glucocorticoid receptor-induced non-muscle caldesmon regulates metastasis in castration-resistant prostate cancer. Oncogenesis 12, 1–12 (2023).

40. S. Jansky, A. K. Sharma, V. Körber, A. Quintero, U. H. Toprak, E. M. Wecht, M. Gartlgruber, A. Greco, E. Chomsky, T. G. P. Grünewald, K. O. Henrich, A. Tanay, C. Herrmann, T. Höfer, F. Westermann, Single-cell transcriptomic analyses provide insights into the developmental origins of neuroblastoma. Nat. Genet. 53, 683–693 (2021).

41. D. R. Rhodes, J. Yu, K. Shanker, N. Deshpande, R. Varambally, D. Ghosh, T. Barrette, A. Pandey, A. M. Chinnaiyan, Large-scale meta-analysis of cancer microarray data identifies common transcriptional profiles of neoplastic transformation and progression. Proc. Natl. Acad. Sci. U. S. A. 101, 9309–9314 (2004).

42. M. T. N. Le, H. Xie, B. Zhou, P. H. Chia, P. Rizk, M. Um, G. Udolph, H. Yang, B. Lim, H. F. Lodish, MicroRNA-125b Promotes Neuronal Differentiation in Human Cells by Repressing Multiple Targets. Mol. Cell. Biol. 29, 5290–5305 (2009).

43. L. Sainero-Alcolado, M. Mushtaq, J. Liaño-Pons, A. Rodriguez-Garcia, Y. Yuan, T. Liu, M. V. Ruiz-Pérez, S. Schlisio, O. Bedoya-Reina, M. Arsenian-Henriksson, Expression and activation of nuclear hormone receptors result in neuronal differentiation and favorable prognosis in neuroblastoma. J. Exp. Clin. Cancer Res. 41 (2022).

44. A. Faisal, L. Vaughan, V. Bavetsias, C. Sun, B. Atrash, S. Avery, Y. Jamin, S. P. Robinson, P. Workman, J. Blagg, F. I. Raynaud, S. A. Eccles, L. Chesler, S. Linardopoulos, The aurora kinase inhibitor CCT137690 downregulates MYCN and sensitizes MYCN-amplified neuroblastoma in Vivo. Mol. Cancer Ther. 10, 2115–2123 (2011).

45. B. Lee, M. T. Muller, SUMOylation enhances DNA methyltransferase 1 activity. Biochem. J. 421, 449–461 (2009).

46. J. S. Kroonen, I. J. de Graaf, S. Kumar, D. F. G. Remst, A. K. Wouters, M. H. M. Heemskerk, A. C. O. Vertegaal, Inhibition of SUMOylation enhances DNA hypomethylating drug efficacy to reduce outgrowth of hematopoietic malignancies. Leukemia 37, 864–876 (2023).

47. N. Braekeveldt, C. Wigerup, D. Gisselsson, S. Mohlin, M. Merselius, S. Beckman, T. Jonson, A. Börjesson, T. Backman, I. Tadeo, A. P. Berbegall, I. Öra, S. Navarro, R. Noguera, S. Påhlman, D. Bexell, Neuroblastoma patient-derived orthotopic xenografts retain metastatic patterns and geno- and phenotypes of patient tumours. Int. J. Cancer 136, E252–E261 (2015).

48. C. U. Persson, K. Von Stedingk, D. Bexell, M. Merselius, N. Braekeveldt, D. Gisselsson, M. Arsenian-Henriksson, S. Påhlman, C. Wigerup, Neuroblastoma patient-derived xenograft cells cultured in stem-cell promoting medium retain tumorigenic and metastatic capacities but differentiate in serum. Sci. Rep. 7, 1–13 (2017).

49. J. Anderson, R. G. Majzner, P. M. Sondel, Immunotherapy of Neuroblastoma: Facts and Hopes. American Association for Cancer Research Inc. (2022). 10.1158/1078-0432.CCR-21-1356.

50. A. Dudek, D. Juric, A. Dowlati, U. Vaishampayan, H. Assad, J. Rodón, B. Chao, B. Wang, J. Gibbs, V. Shinde, S. Friedlander, A. Berger, C. Ward, A. Martinez, R. Gharavi, A. Gomez- Pinillos, I. Proscurshim, A. Olszanski, 476 First-in-human phase 1/2 study of the first-in-class SUMO-activating enzyme inhibitor TAK-981 in patients with advanced or metastatic solid tumors or relapsed/refractory lymphoma: phase 1 results. J. Immunother. Cancer 9, A505– A506 (2021).

51. E. S. Lightcap, P. Yu, S. Grossman, K. Song, M. Khattar, K. Xega, X. He, J. M. Gavin, H. Imaichi, J. J. Garnsey, E. Koenig, H. Zhang, Z. Lu, P. Shah, Y. Fu, M. A. Milhollen, B. A. Hatton, J. Riceberg, V. Shinde, C. Li, J. Minissale, X. Yang, D. England, R. A. Klinghoffer, S. Langston, K. Galvin, G. Shapiro, S. M. Pulukuri, S. Y. Fuchs, D. Huszar, A small-molecule SUMOylation inhibitor activates antitumor immune responses and potentiates immune therapies in preclinical models. Sci. Transl. Med. 13, eaba7791 (2021).

52. T. Kobayashi, K. C. Masoumi, R. Massoumi, Deubiquitinating activity of CYLD is impaired by SUMOylation in neuroblastoma cells. Oncogene 34, 2251–2260 (2015).

53. Q. Chen, R. Deng, X. Zhao, H. Yuan, H. Zhang, J. Dou, R. Chen, H. Jin, Y. Wang, J. Huang, J. Yu, Sumoylation of EphB1 Suppresses Neuroblastoma Tumorigenesis via Inhibiting PKCγ Activation. Cell. Physiol. Biochem. 45, 2283–2292 (2018).

54. Y. Xiang-ming, X. Zhi-qiang, Z. Ting, W. Jian, P. Jian, Y. Li-qun, F. Ming-cui, X. Hong- liang, C. Xu, Z. Yun, SENP1 regulates cell migration and invasion in neuroblastoma. Biotechnol. Appl. Biochem. 63, 435–440 (2016).

55. T. G. F. d. C. Do Nascimento, J. de F. Poloni, M. E. de O. Thomazini, L. R. Cavalli, S. Elifio- Esposito, B. C. Feltes, DNA copy number profiles and systems biology connect chromatin remodeling and DNA repair in high-risk neuroblastoma. Genet. Mol. Biol. 47, e20240007 (2024).

56. W. Yang, W. Paschen, Gene expression and cell growth are modified by silencing SUMO2 and SUMO3 expression. Biochem. Biophys. Res. Commun. 382, 215–218 (2009).

57. W. Yang, J. W. Thompson, Z. Wang, L. Wang, H. Sheng, M. W. Foster, M. A. Moseley, W. Paschen, Analysis of oxygen/glucose-deprivation-induced changes in SUMO3 conjugation using SILAC-based quantitative proteomics. J. Proteome Res. 11, 1108–1117 (2012).

58. Y. J. Lee, J. D. Bernstock, N. Nagaraja, B. Ko, J. M. Hallenbeck, Global SUMOylation facilitates the multimodal neuroprotection afforded by quercetin against the deleterious effects of oxygen/glucose deprivation and the restoration of oxygen/glucose. J. Neurochem. 138, 101– 116 (2016).

59. A. F. Deyrieux, V. G. Wilson, Sumoylation in development and differentiation. Adv. Exp. Med. Biol. 963, 197–214 (2017).

60. L. Gong, W. K. Ji, X. H. Hu, W. F. Hu, X. C. Tang, Z. X. Huang, L. Li, M. Liu, S. H. Xiang, E. Wu, Z. Woodward, Y. Z. Liu, Q. D. Nguyen, D. W. C. Li, Sumoylation differentially regulates Sp1 to control cell differentiation. Proc. Natl. Acad. Sci. U. S. A. 111, 5574–5579 (2014).

61. J. F. Correa-Vázquez, F. Juárez-Vicente, P. García-Gutiérrez, S. V. Barysch, F. Melchior, M. García-Domínguez, The Sumo proteome of proliferating and neuronal-differentiating cells reveals Utf1 among key Sumo targets involved in neurogenesis. Cell Death Dis. 12, 1–14 (2021).

62. B. Mojsa, M. H. Tatham, L. Davidson, M. Liczmanska, E. Branigan, R. T. Hay, Identification of SUMO Targets Associated With the Pluripotent State in Human Stem Cells. Mol. Cell. Proteomics 20, 100164 (2021).

63. O. Wendling, N. B. Ghyselinck, P. Chambon, M. Mark, Roles of retinoic acid receptors in early embryonic morphogenesis and hindbrain patterning. Development 128, 2031–2038 (2001).

64. P. J. McCaffery, J. Adams, M. Maden, E. Rosa-Molinar, Too much of a good thing: Retinoic acid as an endogenous regulator of neural differentiation and exogenous teratogen. John Wiley & Sons, Ltd (2003). 10.1046/j.1460-9568.2003.02765.x.

65. V. Rodriguez, R. Bailey, M. Larion, M. R. Gilbert, Retinoid receptor turnover mediated by sumoylation, ubiquitination and the valosin-containing protein is disrupted in glioblastoma. Sci. Rep. 9, 1–13 (2019).

66. J. S. Lim, S. H. Park, K. L. Jang, All-trans retinoic acid induces cellular senescence by up- regulating levels of p16 and p21 via promoter hypomethylation. Biochem. Biophys. Res. Commun. 412, 500–505 (2011).

67. I. Westerlund, Y. Shi, K. Toskas, S. M. Fell, S. Li, O. Surova, E. Södersten, P. Kogner, U. Nyman, S. Schlisio, J. Holmberg, Combined epigenetic and differentiation-based treatment inhibits neuroblastoma tumor growth and links HIF2α to tumor suppression. Proc. Natl. Acad. Sci. U. S. A. 114, E6137–E6146 (2017).

68. F. Juarez-Vicente, N. Luna-Pelaez, M. Garcia-Dominguez, The Sumo protease Senp7 is required for proper neuronal differentiation. Biochim. Biophys. Acta - Mol. Cell Res. 1863, 1490–1498 (2016).

69. E. Gonçalves, R. C. Poulos, Z. Cai, S. Barthorpe, S. S. Manda, N. Lucas, A. Beck, D. Bucio- Noble, M. Dausmann, C. Hall, M. Hecker, J. Koh, H. Lightfoot, S. Mahboob, I. Mali, J. Morris, L. Richardson, A. J. Seneviratne, R. Shepherd, E. Sykes, F. Thomas, S. Valentini, S. G. Williams, Y. Wu, D. Xavier, K. L. MacKenzie, P. G. Hains, B. Tully, P. J. Robinson, Q. Zhong, M. J. Garnett, R. R. Reddel, Pan-cancer proteomic map of 949 human cell lines. Cancer Cell 40, 835–849.e8 (2022).

