## Supplementary figures for "SUMOylation is a Therapeutic Vulnerability in High-risk Neuroblastoma"

A

| SUMOylation-regulating factor | Kocac cohort (n=476) | SECQ cohort (n=498) | Cangelosi cohort (n=419) | Versteeg cohort (n=88) |
| --- | --- | --- | --- | --- |
| **SAE1** | **** (p=5.8e-08) | **** (p=4.8e-15) | **** (p=1.2e-08) | **** (p=3.6e-05) |
| **SAE2 (UBA2)** | **** (p=3.9e-07) | **** (p=2.4e-17) | ** (p=5.0e-03) | **** (p=7.2e-06) |
| **UBE2I** | **** (p=1.7e-07) | **** (p=7.2e-10) | **** (p=3.8e-09) | *** (p=4.1e-04) |
| SUMO1 | ** (p=3.3e-03) | **** (p=2.9e-06) | Ns. (p=0.457) | Ns. (p=0.547) |
| SUMO2 | Ns. (p=0.081) | **** (p=3.3e-08) | N/A | N/A |
| SUMO3 | Ns. (p=0.506) | *** (p=1.8e-04) | Ns. (p=0.421) | Ns. (p=0.355) |
| PIAS1 | * (p=0.011) | Ns. (p=0.730) | Ns. (p=0.244) | Ns. (p=0.816) |
| PIAS2 | *** (p=1.1e-04) | ** (p=6.0e-03) | ** (p=7.4e-03) | Ns. (p=0.430) |
| PIAS3 | **** (p=1.5e-06) | **** (p=4.2e-11) | Ns. (p=0.117) | Ns. (p=0.518) |
| PIAS4 | * (p=0.011) | Ns. (p=0.069) | * (p=0.010) | Ns. (p=0.282) |
| SEN1 | *** (p=6.4e-04) | * (p=0.016) | Ns. (p=0.353) | Ns. (p=0.401) |
| **SEN2** | **** (p=7.0e-10) | **** (p=4.3e-05) | **** (p=1.6e-07) | ** (p=2.7e-03) |
| SEN3 | Ns. (p=0.388) | Ns. (p=0.218) | Ns. (p=0.503) | Ns. (p=0.902) |
| SEN5 | *** (p=2.9e-04) | **** (p=6.7e-07) | Ns. (p=0.089) | Ns. (p=0.053) |
| **SEN6** | ** (p=7.2e-03) | **** (p=1.3e-06) | **** (p=3.5e-06) | **** (p=2.1e-06) |
| **SEN7** | **** (p=4.8e-11) | **** (p=1.7e-10) | **** (p=1.8e-14) | **** (p=9.8e-07) |
| PML | Ns. (p=0.268) | *** (p=3.1e-04) | Ns. (p=0.766) | Ns. (p=0.834) |
| RANBP2 | Ns. (p=0.716) | * (p=0.014) | * (p=0.044) | Ns. (p=0.509) |
| PC2 (CBX4) | Ns. (p=0.053) | Ns. (p=0.708) | **** (p=6.4e-06) | * (p=0.018) |
| ZNF451 | Ns. (p=0.305) | Ns. (p=0.257) | **** (p=4.0e-06) | ** (p=1.3e-03) |
| RSUME (RWDD3) | Ns. (p=0.222) | Ns. (p=0.284) | Ns. (p=0.095) | Ns. (p=0.074) |

B

| SUMOylation-regulating factor | Alteration frequency in patients (%; n=59) |  | Alteration frequency in cell lines (%; n=17) |  |
| --- | --- | --- | --- | --- |
|  | Amplification | Deletion | Amplification | Deletion |
| SAE1 | - | - | - | 11.8 |
| SAE2 | - | - | - | - |
| UBE2I | 1.7 | - | - | - |
| SUMO1 | - | 1.7 | - | - |
| SUMO2 | 8.5 | - | 29.4 | - |
| SUMO3 | - | - | - | - |
| PIAS1 | - | - | - | - |
| PIAS2 | 1.7 | - | - | 23.5 |
| PIAS3 | - | - | 11.8 | - |
| PIAS4 | - | - | - | - |
| SEN1 | - | - | - | - |
| SEN2 | - | - | - | - |
| SEN3 | - | - | - | 23.5 |
| SEN5 | - | - | - | - |
| SEN6 | - | - | - | - |
| SEN7 | - | - | - | - |
| PML | - | - | - | - |
| RANBP2 | - | - | - | 11.8 |
| PC2 | 10.2 | - | 35.3 | - |
| ZNF451 | - | - | - | - |
| RSUME | - | 1.7 | - | 17.3 |

C

| MYCN status | Cell line | Genetic alterations of SUMOylation regulators |
| --- | --- | --- |
| Non-amplified | SK-N-AS | Amplification: PC2, SUMO2<br>Deletion: PIAS2, SENP3 |
|  | SH-SY5Y | Mutation: SAE1 (K45E) |
|  | SK-N-SH* | Mutation: SAE1 (K45E) |
|  | SK-N-FI | No alterations |
| MYCN-amplified | CHP-126 | Amplification: PC2, SUMO2 |
|  | CHP-212 | Amplification: PC2, SUMO2<br>Deletion: RSUME |
|  | IMR-32 | Amplification: PC2, SUMO2 |
|  | KELLY | Deletion: RANBP2 |
|  | KP-N-RT-BM-1 | Deletion: RSUME |
|  | KP-N-SI9s | Amplification: PIAS3<br>Deletion: PIAS2 |
|  | KP-N-YN | Deletion: SAE1, SENP3 |
|  | MHH-NB-11 | No alterations |
|  | NB-1 | Amplification: PIAS3<br>Deletion: PIAS2 |
|  | NH-6 | Amplification: PC2<br>Deletion: SENP3 |
|  | SIMA | Amplification: PC2, SUMO2<br>Deletion: RANBP2 |
|  | SK-N-BE(2) | Deletion: PIAS2, SENP3, RSUME |
|  | SK-N-DZ | Deletion: SAE1 |

**Supplementary Figure S1. SUMO pathway components have prognostic significance and copy number alterations in NB. (A)** Association between the mRNA expression levels of SUMO machinery components and OS in NB patients from the Kocak, SEQC, Cangelosi, and Versteeg cohorts. Bold text denotes SUMO machinery components with prognostic significance in all analyzed cohorts. Median mRNA expression value was used as a cutoff for group stratification. Red/blue = high/low expression correlates with poor prognosis as determined with the log-rank test (\* $p < 0.05$, \*\* $p < 0.01$, \*\*\* $p < 0.001$, \*\*\*\* $p < 0.0001$). **(B)** Frequency of copy number alterations of SUMOylation-regulating factors in NB patients and cell lines analyzed from the TARGET cohort and CCLE. Amplification and deletion status were inferred from the GISTIC algorithm. **(C)** Copy number alterations and mutations of SUMOylation-regulating factors in *MYCN*-non-amplified and *MYCN*-amplified NB cell lines analyzed from the CCLE.

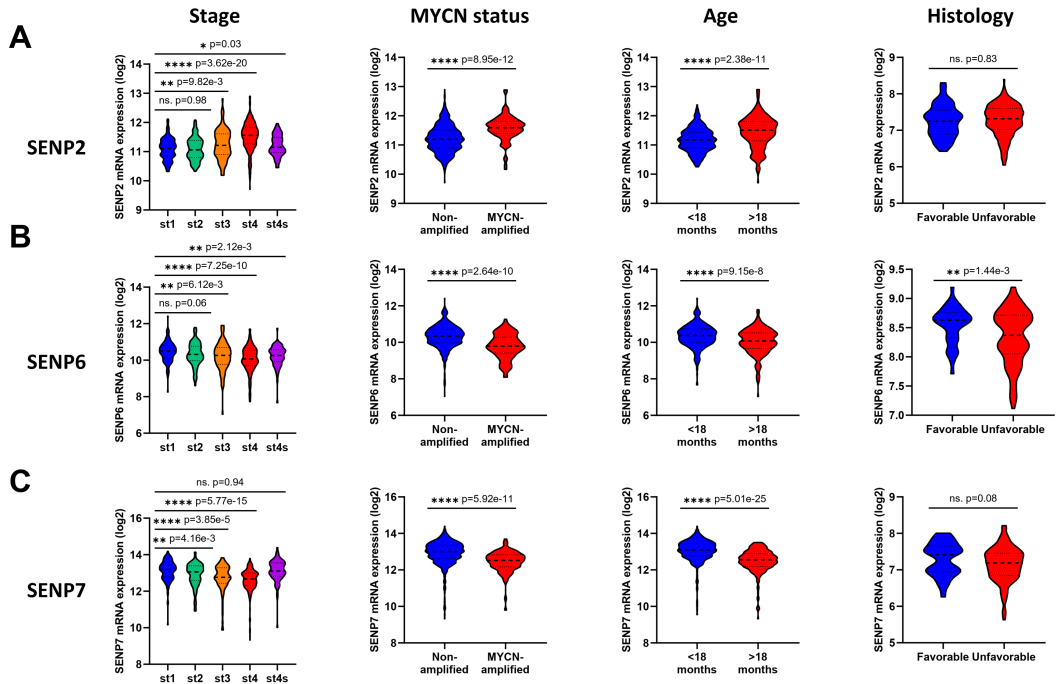

**D**

| SUMOylation-regulating factor | Stage (st. 1 vs. 4) | MYCN status (non-amp vs. MYCN-amp) | Age (<18 vs. >18 months) | Histology (favourable vs. unfavourable) |
| --- | --- | --- | --- | --- |
| SUMO1 | ** (p=8.89e-03) | **** (p=4.85e-07) | Ns. (p=0.925) | Ns. (p=0.759) |
| SUMO2 | ** (p=4.93e-03) | Ns. (p=0.386) | Ns. (p=0.440) | Ns. (p=0.0508) |
| SUMO3 | Ns. (p=0.180) | *** (p=3.23e-04) | Ns. (p=0.139) | Ns. (p=0.968) |
| PIAS1 | Ns. (p=0.111) | * (p=0.020) | **** (p=5.35e-06) | Ns. (p=0.401) |
| PIAS2 | **** (p=3.93e-05) | **** (p=2.74e-05) | **** (p=8.29e-19) | * (p=0.045) |
| PIAS3 | **** (p=4.98e-18) | **** (p=2.95e-17) | ** (p=5.92e-03) | **** (p=1.37e-5) |
| PIAS4 | **** (p=1.03e-09) | Ns. (p=0.432) | * (p=0.026) | Ns. (p=0.889) |
| SEN1 | Ns. (p=0.055) | ** (p=1.18e-03) | **** (p=8.19e-05) | Ns. (p=0.0760) |
| SEN3 | Ns. (p=0.412) | Ns. (p=0.773) | Ns. (p=0.408) | Ns. (p=0.323) |
| SEN5 | *** (p=9.19e-04) | * (p=0.028) | *** (p=4.15e-04) | Ns. (p=0.078) |
| PML | * (p=0.029) | * (p=0.037) | Ns. (p=0.474) | Ns. (p=0.185) |
| RanBP2 | **** (p=1.93e-05) | Ns. (p=0.730) | ** (p=9.90e-03) | Ns. (p=0.410) |
| PC2 (CBX4) | *** (p=1.02e-04) | Ns. (p=0.444) | **** (p=9.87e-09) | Ns. (p=0.363) |
| ZNF451 | **** (p=8.15e-10) | Ns. (p=0.318) | **** (p=2.32e-09) | Ns. (p=0.347) |
| RSUME (RWDD3) | Ns. (p=0.180) | *** (p=4.99e-04) | Ns. (p=0.146) | Ns. (p=0.160) |

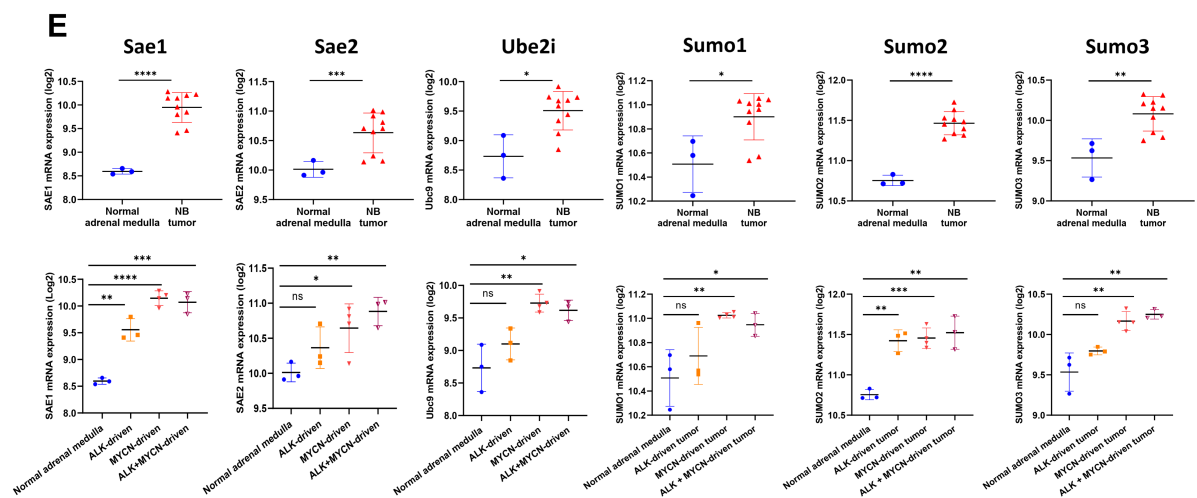

**Supplementary Figure S2. Additional analyses of the mRNA expression of SUMO machinery components in NB patients and NB mouse models.** Violin plots depicting the association of **(A)** *SENP2*, **(B)** *SENP6*, and **(C)** *SENP7* expression with low- and high-risk clinical features in NB patients. Correlation with disease stage (st) was analyzed from patients with st1 (n=153), st2 (n=113), st3 (n=91), st4 (n=214), and st4s (n=78) tumors. Association with *MYCN* amplification status was analyzed between *MYCN*-non-amplified (n=550) and *MYCN*-amplified (n=93) tumors. Correlation with diagnosis age was analyzed from patients who were <18 months (n=414) and >18 months (n=235) of age. Correlation with histopathology was evaluated from tumors with favorable (n=37) and unfavorable (n=183) histology. Association with stage, *MYCN*, and age were analyzed from the Kocak cohort and correlation with histology was analyzed from the TARGET cohort. Welch's t-test was used for the statistical testing (\* $p < 0.05$, \*\* $p < 0.01$, \*\*\* $p < 0.001$, \*\*\*\* $p < 0.0001$). **(D)** Table of clinical associations of SUMO machinery components that did not have prognostic impact on all of the cohorts. Association of mRNA expression with stage, *MYCN*, and age were analyzed from the Kocak cohort and correlation with histology was analyzed from the TARGET cohort. Red indicates higher expression in high-risk (stage 4, *MYCN*-amplified, <18 months, and unfavorable histology) group and blue indicates higher expression in low-risk (stage 1, *MYCN*-non-amplified, >18 months, and favorable histology) group. Welch's t-test was used for the statistical testing (\* $p < 0.05$, \*\* $p < 0.01$, \*\*\* $p < 0.001$, \*\*\*\* $p < 0.0001$). **(E)** Scatter dot plots with SD of the expression of *Sae1*, *Sae2*, *Ube2i*, and *Sumo1–3* in mouse normal adrenal medulla (n=3), NB tumor (n=10), and different tumor subtypes consisting of *ALK*- (n=3), *MYCN*- (n=4), and *ALK*+*MYCN*-driven (n=3) tumors of NB murine models. Analysis was done from the GSE32386 dataset. Welch's t-test was used for statistical testing (\* $p < 0.05$, \*\* $p < 0.01$, \*\*\* $p < 0.001$, \*\*\*\* $p < 0.0001$).

A

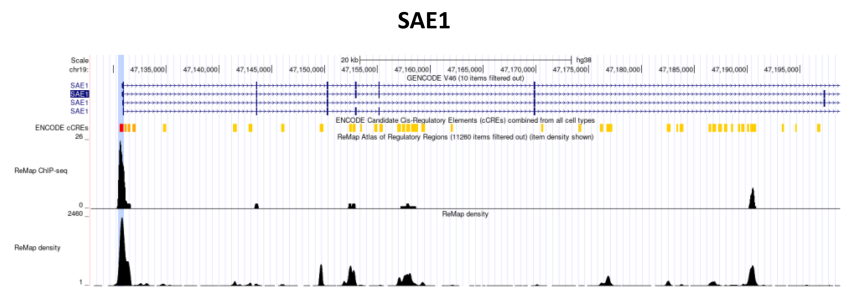

B

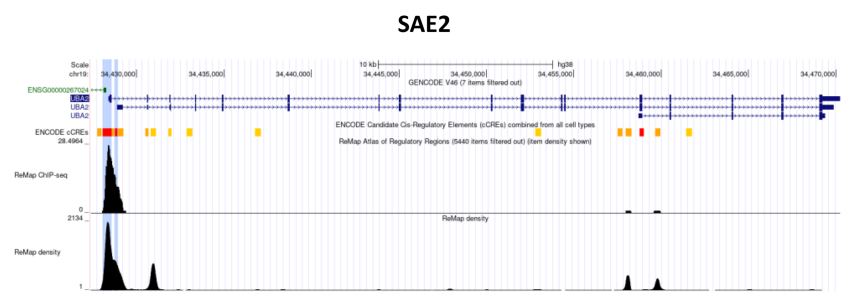

C

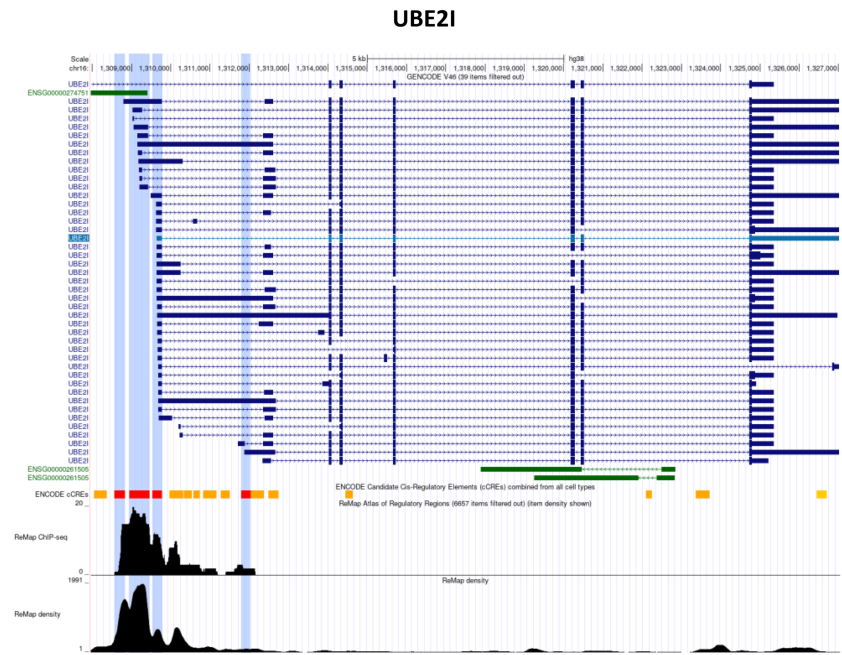

D

Overall survival in MYCN-non-amplified NB patients

| SUMOylation-regulating factor | Kocak cohort (n=405) | SECQ cohort (n=401) | Cangelosi cohort (n=334) | Versteeg cohort (n=72) |
| --- | --- | --- | --- | --- |
| SAE1 | *** (p=5.90e-04) | **** (p=1.21e-08) | **** (p=5.99e-05) | *** (p=4.10e-04) |
| SAE2 | * (p=0.012) | **** (p=1.54e-08) | * (p=0.039) | * (p=0.035) |
| UBE2I | * (p=0.049) | ** (p=1.75e-03) | **** (p=9.96e-05) | *** (p=5.02e-04) |

**Supplementary Figure S3. *SAE1*, *SAE2*, and *UBE2I* are putative transcriptional targets of MYCN.** Visualization of ReMap ChIP-seq tracks for MYCN in **(A)** *SAE1*, **(B)** *SAE2*, and **(C)** *UBE2I* with ENCODE candidate promoter-like signatures marked in red and enhancer-like signatures in yellow. Highlighting in blue indicate peaks for MYCN on the promoter-like signatures of ReMap ChIP-seq data pooled from all available ENCODE data. The graphs were generated with the UCSC Genome Browser. **(D)** Impact of *SAE1*, *SAE2*, and *UBE2I* on OS MYCN-non-amplified NB patients analyzed from the Kocak, SEQC, Cangelosi, and Versteeg cohorts. Median mRNA expression value was used as a cutoff for group stratification. Red = high expression correlates with poor prognosis as determined with the log-rank test (\* $p < 0.05$, \*\* $p < 0.01$, \*\*\* $p < 0.001$, \*\*\*\* $p < 0.0001$).

### SAE1

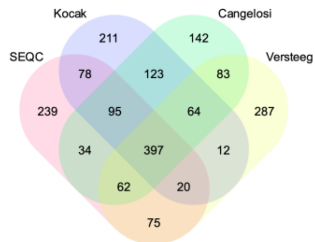

### GO:BP gene sets

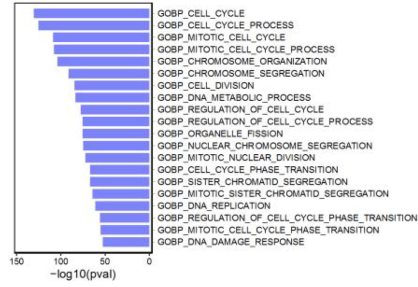

### Hallmark gene sets

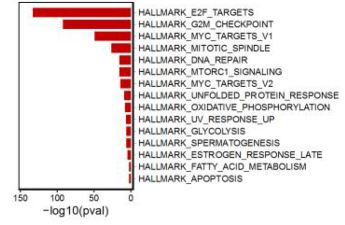

### SAE2

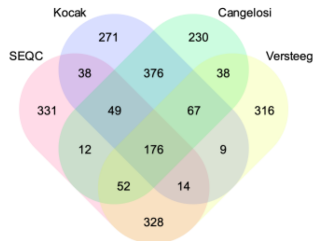

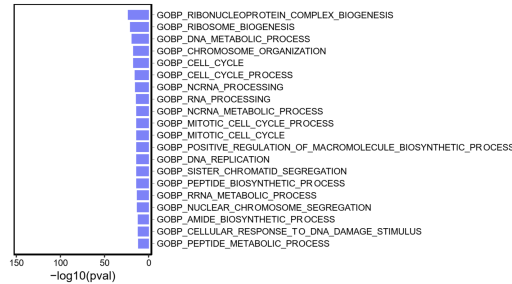

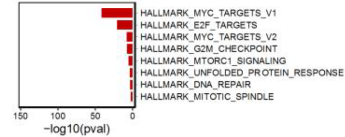

### UBE2I

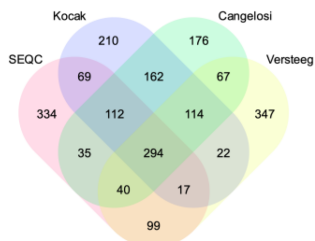

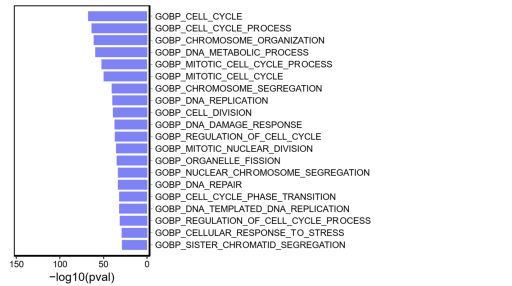

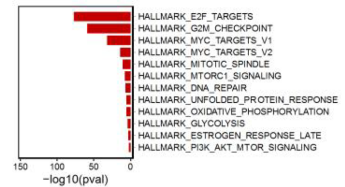

### SUMO1

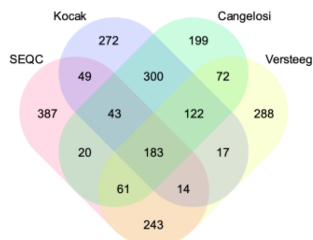

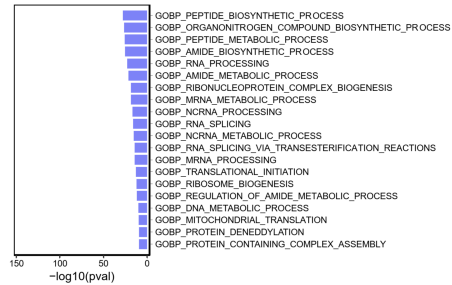

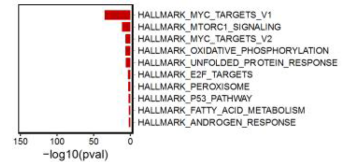

### SUMO2

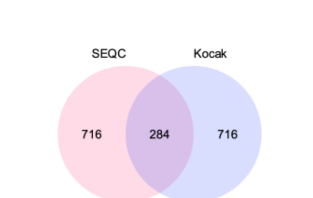

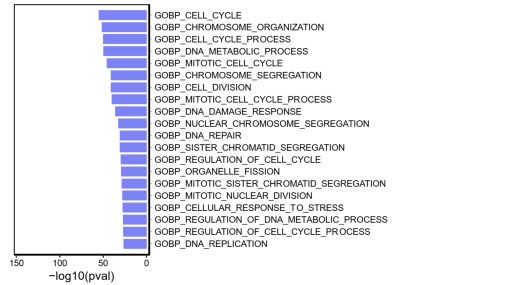

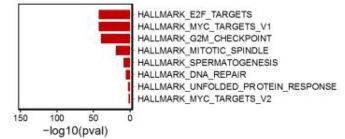

### SUMO3

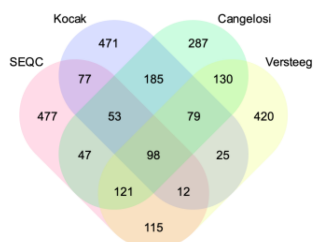

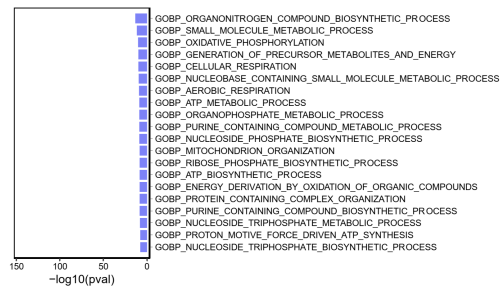

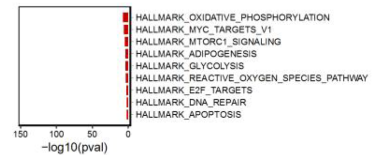

**Supplementary Figure S4. SUMOylation-promoting factors correlate strongly with cell cycle related gene sets.** Venn diagrams representing the number of commonly co-expressed genes for *SAE1*, *SAE2*, *UBE2I*, *SUMO1*, *SUMO2*, and *SUMO3* and bar graphs depicting the top 10 most significantly associated GO:BP and Hallmark gene sets. The lists of commonly co-expressed genes were identified from the top 1000 positively co-expressed genes in Kocak, SEQC, Cangelosi, and Versteeg cohorts. P values ($-\log_{10}$) derived from hypergeometric distribution were used for representing the statistical significance of gene sets.

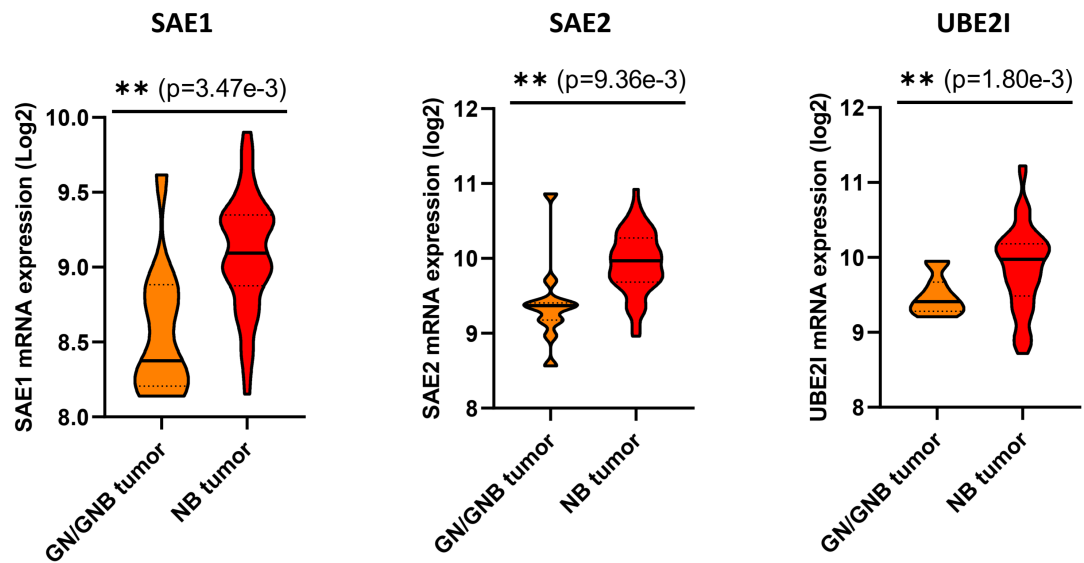

**Supplementary Figure S5. Expression levels of SUMOylation-promoting factors are higher in NB compared to other neuroblastic tumor types.** Violin plots comparing the expression levels of *SAE1*, *SAE2*, and *UBE2I* between ganglioneuroma/ganglioneuroblastoma (GN/GNB) tumor (n=11) and NB tumor (n=53) samples. Data was analyzed from GSE12460. Welch's t-test was used for statistical testing (\*\* $p < 0.01$).

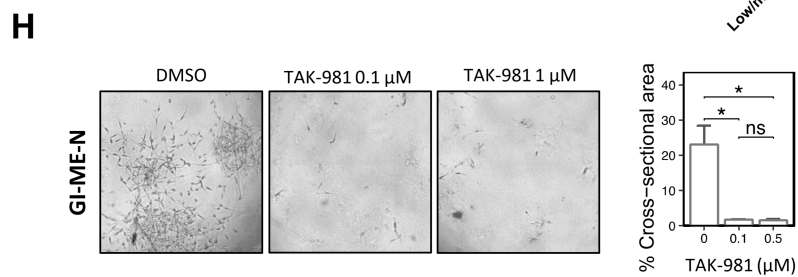

**Supplementary Figure S6. NB cell lines are sensitive to *SAE1* and *SAE2* knockout irrespective of *MYCN* or *ALK* status.** (A) DepMap Oncotree lineages ranked based on the mean gene effect of *UBE2I* analyzed from the CRISPR (DepMap Public 26Q1+Score, Chronos) dataset (n=1208). The blue line indicates the mean gene effect across all cell lines. (B) DepMap Oncotree primary diseases ranked based on the mean gene effect of *SAE1* and *SAE2*; only primary diseases where the gene effect differed statistically (Welch's t-test) from the mean gene effect of all other cell lines are shown in the graphs. The blue line indicates the mean gene effect of across all cell lines. (C) Combined mean gene effect of *SAE1* and *SAE2* in *MYCN*-non-amplified (n=10) and *MYCN*-amplified (n=31) NB cell lines tested with Student's t-test. (D) *SAE1* gene effect, *SAE2* gene effect, and the combined mean gene effect of *SAE1* and *SAE2* in wild type *ALK* (n=30) and *ALK*-mutated/amplified (n=11) NB cell lines tested with Student's t-test. (E) Late neuroblast signature (LNS) score in NB cell lines available in DepMap. (F) Combined mean gene effect of *SAE1* and *SAE2* in NB cell lines expressing high (top tertile) and low/medium (bottom tertiles) late neuroblast score tested with Student's t-test. (G) Comparison of TAK-981 IC50 values (-log, M) in wild type *ALK* (n=4) and *ALK*-mutated/amplified (n=4) NB cell lines tested with Student's t-test. (H) Representative brightfield images of *MYCN*-non-amplified GI-ME-N cells treated with DMSO and TAK-981 (0.1 and 0.5 μM) grown in Matrigel (day 10). Bar graph with SEM indicates the % of cross-sectional area; data was analyzed from four brightfield images/treatment group. Student's t-test was used for statistical testing (\* p < 0.05).

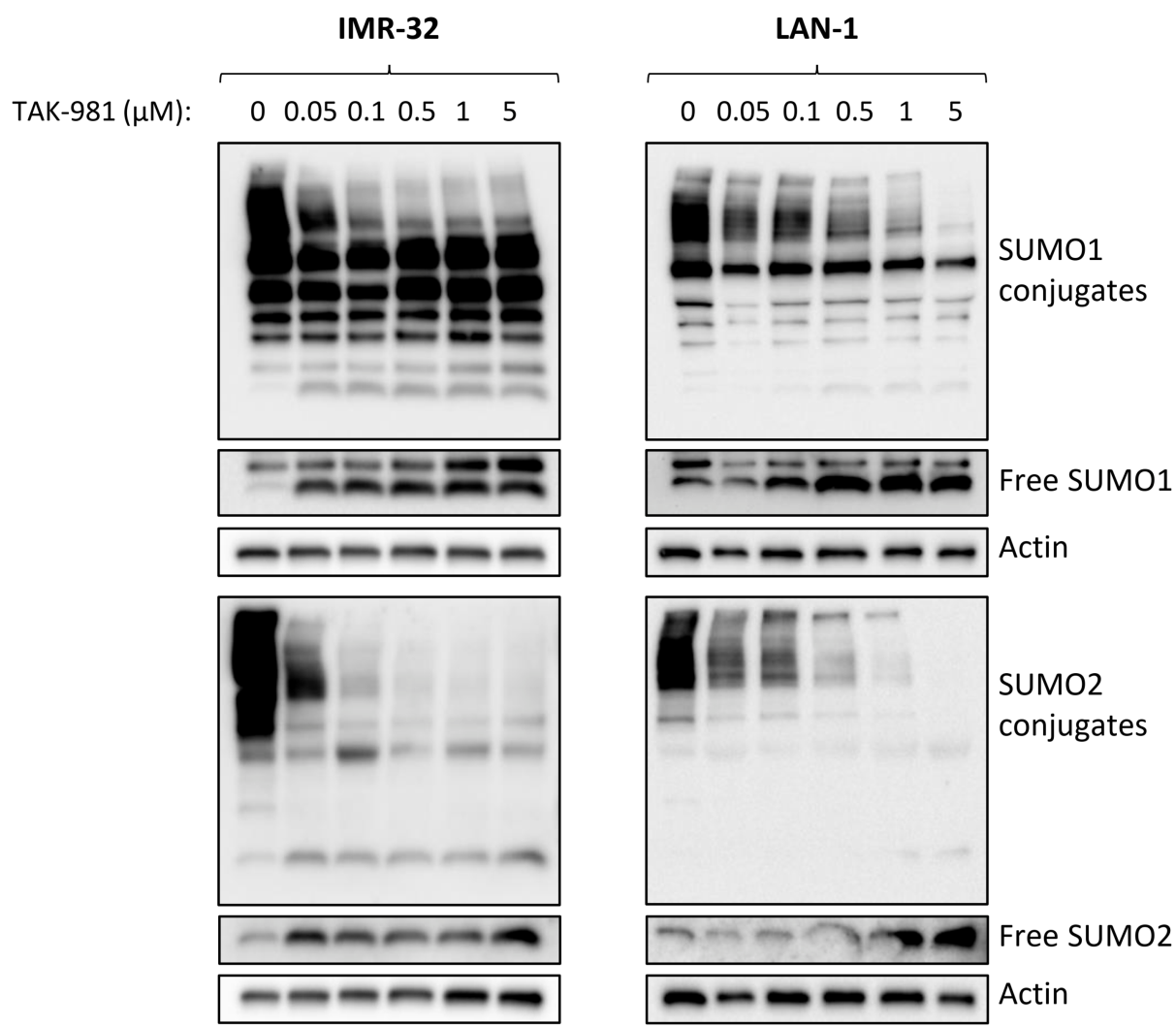

**Supplementary Figure S7. TAK-981 inhibits the level of global SUMOylation in NB cells.**

Representative western blots of the conjugation levels of SUMO1 and SUMO2 in IMR-32 and LAN-1 cells treated with increasing concentrations of TAK-981 for 48 hours. Longer exposures of SUMO1 and SUMO2 are shown below to illustrate the changes in the levels of free SUMO1 and SUMO2.

**A**

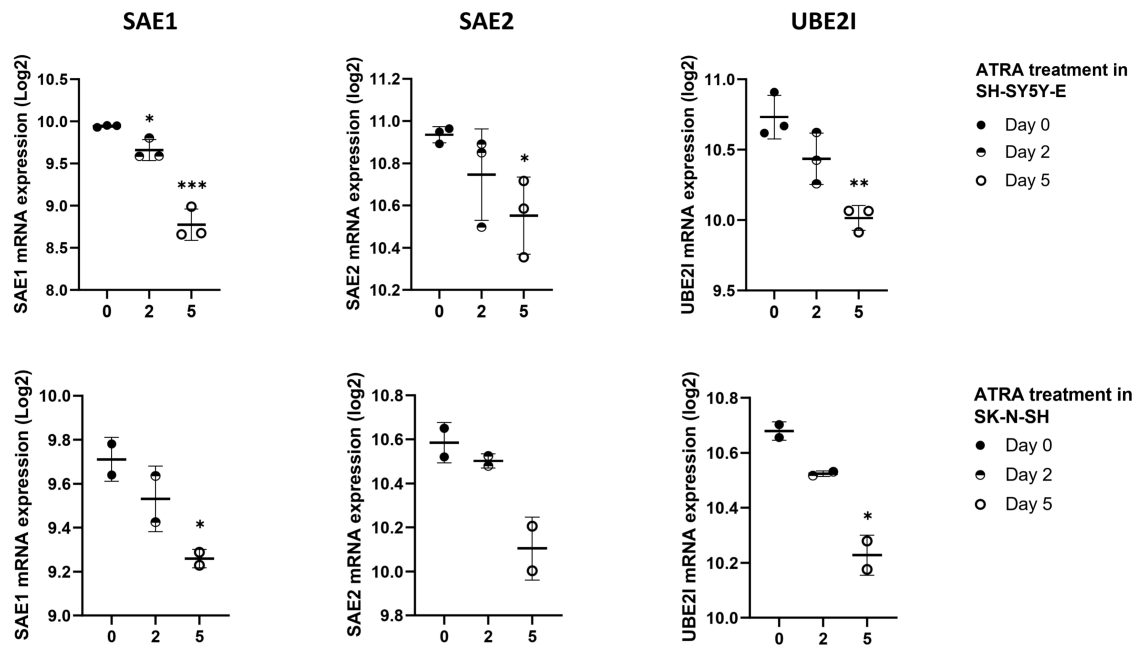

**B**

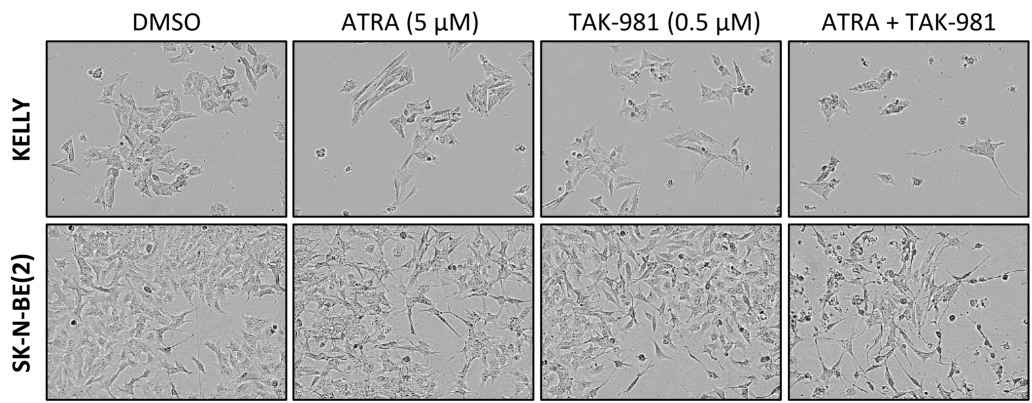

**Supplementary Figure S8. The effect of ATRA on the mRNA expression of SUMOylation-promoting factors in an additional dataset and morphology images of KELLY and SK-N-BE(2) cells after combination treatment with ATRA and TAK-981.** (A) Expression of *SAE1*, *SAE2*, and *UBE2I* are represented as scatter dot plots with SD after 0, 2, and 5 days of ATRA treatment (5 μM) in SK-N-SH and SH-SY5Y-E cells. Data was analyzed from the GSE9169 dataset. Statistical significance was determined with Student's t-test (\* $p < 0.05$, \*\* $p < 0.01$, \*\*\* $p < 0.001$). (B) Representative brightfield images of cell morphology of KELLY and SK-N-BE(2) cells treated with ATRA (5 μM), TAK-981 (0.5 μM) and the combination of ATRA and TAK-981 taken with IncuCyte after 4 days of treatment.

A

TAK-981 + Tozasertib combination in IMR-32 cells (3D)

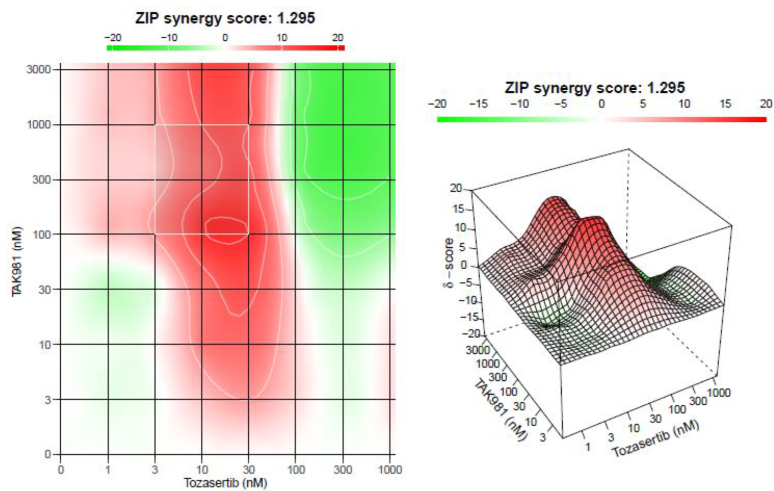

TAK-981 + Alisertib combination in IMR-32 cells (3D)

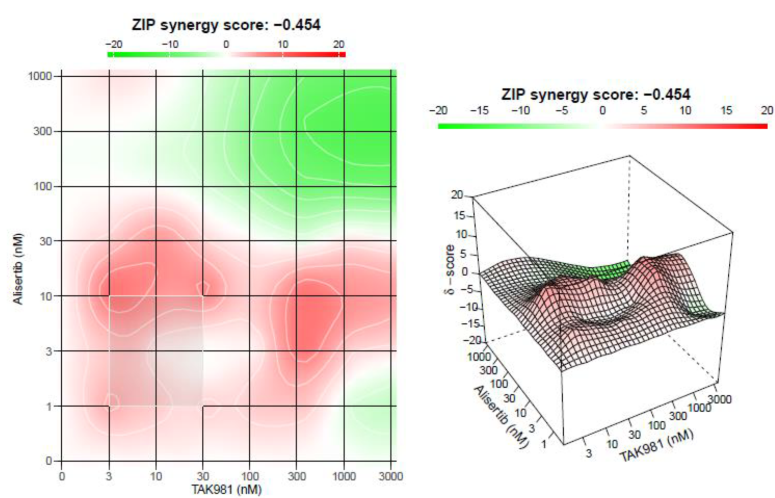

B

TAK-981 + Decitabine combination in IMR-32 cells (3D)

**Supplementary Figure S9. TAK-981 combination treatments with pan-aurora kinase inhibitor tozasertib, selective aurora A inhibitor alisertib, and DNMT inhibitor decitabine. (A)** Drug synergy scores of the TAK-981 + tozasertib and TAK-981 + alisertib combinations. IMR-32 cells were grown in 3D spheroid conditions, and cells were exposed to concentration matrices of the indicated drugs for 72 hours. Drug interactions were evaluated using the ZIP synergy scoring model. **(B)** Dose-response curves of decitabine and TAK-981, and dose-response matrix of the TAK-981 + decitabine combination after 72 hours in IMR-32 spheroids.
