## Supplementary Table S1 for "SUMOylation is a Therapeutic Vulnerability in High-risk Neuroblastoma"

**Table 1. List of antibodies used for western blot analysis.** Abbreviations: mAb monoclonal antibody, pAb polyclonal antibody.

| **Target protein** | **Catalog #** | **Manufacturer** | **Clonality** | **Dilution** |
| --- | --- | --- | --- | --- |
| **SAE1** | ab185552 | Abcam | Rabbit mAb | 1:1000 |
| **SAE2** | ab185955 | Abcam | Rabbit mAb | 1:1000 |
| **UBC9** | ab75854 | Abcam | Rabbit mAb | 1:1000 |
| **SUMO1** | ab133352 | Abcam | Rabbit mAb | 1:1000 |
| **SUMO2** | ab212838 | Abcam | Mouse mAb | 1:750 |
| **RARα** | ab275745 | Abcam | Rabbit mAb | 1:1000 |
| **PC2** | #14013 | Cell Signaling Technology | Rabbit mAb | 1:1000 |
| **pAKT (s473)** | #4060 | Cell Signaling Technology | Rabbit mAb | 1:1000 |
| **AKT** | #4691 | Cell Signaling Technology | Rabbit mAb | 1:1000 |
| **Cleaved PARP** | #5625 | Cell Signaling Technology | Rabbit mAb | 1:1000 |
| **Cleaved caspase-3** | #9664 | Cell Signaling Technology | Rabbit mAb | 1:750 |
| **p27** | #3686 | Cell Signaling Technology | Rabbit mAb | 1:1000 |
| **NTRK1** | #2510 | Cell Signaling Technology | Rabbit mAb | 1:750 |
| **PHOX2B** | #83811 | Cell Signaling Technology | Rabbit pAb | 1:1000 |
| **MYCN** | #84406 | Cell Signaling Technology | Rabbit mAb | 1:1000 |
| **DNMT1** | #5032 | Cell Signaling Technology | Rabbit mAb | 1:1000 |
| **γH2AX** | ##9718 | Cell Signaling Technology | Rabbit mAb | 1:1000 |
| **Actin** | sc-47778 | Santa Cruz Biotechnology | Mouse mAb | 1:1000 |
| **Vinculin** | sc-73614 | Santa Cruz Biotechnology | Mouse mAb | 1:1000 |
